# Selective augmentation of intestinal immunity by CD22-dependent SHP-1 control of β_7_ integrin expression

**DOI:** 10.1101/2020.01.31.929687

**Authors:** Romain Ballet, Carolin Brandl, Ningguo Feng, Jeremy Berri, Julian Cheng, Borja Ocón, Amin Alborzian Deh Sheikh, Alex Marki, Clare L. Abram, Clifford A. Lowell, Takeshi Tsubata, Harry B. Greenberg, Matthew S. Macauley, Klaus Ley, Lars Nitschke, Eugene C. Butcher

## Abstract

The regulation of integrin expression and function controls interactions of immune cells and targets their trafficking locally and systemically. We show here that the tyrosine phosphatase SHP-1 is required for lymphocyte surface expression of the intestinal immune response-associated integrin β_7_, but not for β_1_ or β_2_ integrins. *Viable motheaten* mice deficient for SHP-1 have less β_7_ on T cells and lack β_7_ on B cells. SHP-1 function is targeted in B cells by the B cell specific lectin CD22 (Siglec-2), suggesting a potential role for CD22 in β_7_ expression. CD22-deficiency on B cells phenocopies the effects of SHP-1 haplodeficiency. Mechanistically, we show that SHP-1 suppresses β_7_ endocytosis: internalization of β_7_ but not β_1_ integrin is accelerated in SHP-1^+/−^ and CD22^−/−^ B cells. Moreover, mutations in CD22 cytoplasmic SHP1-binding ITIM sequences reduce α_4_β_7_ comparably, and loss of CD22 lectin activity has an intermediate effect suggesting a model in which the CD22 ITIM sequences recruit SHP-1 to control β_7_ expression. Integrin α_4_β_7_ selectively contributes to cell interactions in intestinal immunity. Consistent with this, CD22 deficient and SHP-1^+/−^ B cells display reduced β_7_-dependent homing to gut associated Peyer’s patches (PP); and CD22-deficiency impairs intestinal but not systemic antibody responses and delays clearance of the gut pathogen rotavirus. The results define a novel role for SHP-1 in the differential control of leukocyte integrins and an unexpected integrin β_7_-specific role for CD22-SHP-1 interplay in mucosal immunity.

## Introduction

Integrins form a large family of cell surface heterodimeric receptors involved in cell-cell or cell-ECM (extra cellular matrix) interactions. Each heterodimer comprises an α-subunit and β-subunit with unique binding properties. By targeting leukocytes to ligand-expressing tissues, integrins are essential for immune homeostasis and inflammation. For instance, the integrin α_4_β_7_ functions as a B and T cell homing receptor for the mucosal vascular addressin (MAdCAM-1) expressed in the gut-associated lymphoid tissues (GALT) (Bargatze et al., 1995; Berlin et al., 1993). Peyer’s patches (PP) and GALT are major sites of B cell activation and humoral immune induction. Activated B cells undergo isotype class switching in PP and differentiate into migratory IgA secreting plasmablasts that home to mucosal surfaces (Butcher et al., 2005; Kraal et al., 1982), where local production of secretory IgA provides immune protection. Perhaps in support of this role, B cells predominate in murine PP, contrasting with peripheral lymph nodes (PLN) where T cells are the majority (Stevens et al., 1982). This homing preference correlates with higher levels of α_4_β_7_ on B cells than on T cells (Tang et al., 1998), but the mechanisms responsible for differential α_4_β_7_ expression and its essential role in enhanced B cell recruitment to PP have not been explored.

Fine-tuning of integrin activation is key to integrin function. Aberrant activation of integrin often inhibits, rather than promotes, leukocyte trafficking to target tissues, as shown for β_7_ and lymphocytes homing to the gut (Eun et al., 2007), or β_2_ and lymphocytes homing to the PLN (Park et al., 2010). Molecules and signaling pathways involved in the regulation of integrin activation are now well understood while the mechanisms controlling integrin expression have been less explored. Amongst the negative regulators of integrin activation is the tyrosine phosphatase SHP-1 (Src homology region 2 domain-containing phosphatase-1) found in all hematopoietic cells (Neel et al., 2003; Zhang et al., 2000). SHP-1 controls the deactivation of the LFA-1 integrin (α_L_β_2_) to prevent aberrant adhesion of leukocytes to β_2_ integrin ligands (Kruger et al., 2000; Roach et al., 1998; Stadtmann et al., 2015), or T cell adhesion to antigen presenting cells (Sauer et al., 2016). SHP-1 binds phosphorylated ITIM (Immunoreceptor tyrosine-based inhibitory motif) sequences, often acting in conjunction with ITIM-bearing molecules to inhibit downstream signaling responses. In B cells, upon antigen stimulation, the phosphorylated ITIM sequences of CD22 (Sialic acid-binding Ig-like lectin 2 or Siglec-2) recruits SHP-1 to inhibit downstream components of the BCR (B cell receptor)-induced Ca^2+^ signaling (Blasioli et al., 1999; Campbell and Klinman, 1995; Law et al., 1996).

Here we report a new and unexpected function for SHP-1 with profound and organotypic effects on mucosal immune responses. We show that SHP-1 acts cell intrinsically to maintain α_4_β_7_ surface expression by inhibiting β_7_ endocytosis in lymphocytes without impacting other integrins. We show that haploinsufficiency for SHP-1 causes severe impairment of B and T lymphocyte homing to the PP. In B cells, we identified CD22 as being the ITIM-bearing molecule recruiting SHP-1 to the cell surface to mediate the effects on β_7_ regulation. B cells lacking CD22, or expressing CD22 with mutated SHP-1-binding ITIM domains have similar reductions in β_7_ expression and in PP homing. Using CD22-deficiency as a surrogate of the SHP-1 deficiency in B cells, we provide evidence that lack of CD22 reduces antigen specific antibody responses after oral but not nasal or intramuscular immunization. Applied to a mouse model of rotavirus infection, CD22-deficient animals showed a reduced protective immune response in the gut. Our findings highlight a novel and selective SHP-1-dependent ITIM-mediated mechanism of β_7_ integrin regulation by CD22 in B cells, and demonstrate its importance to efficient intestinal antibody responses.

## Results

### SHP-1 selectively controls α_4_β_7_ expression in lymphocytes

The role of SHP-1 in integrin signaling (activation) has been previously addressed without a thorough analysis of integrin homeostasis. We used the *viable motheaten* mutant mice (me^V^/ me^V^, or SHP-1^−/−^) as a model of SHP-1 deficiency to address the question of integrin expression in T and B cells. Immunofluorescent staining with antibodies to the α_4_β_7_ heterodimer or to the β_7_ subunit revealed a dramatic reduction in the median fluorescence intensity (MFI) on SHP-1-deficient splenic B cells as compared to wild-type controls (>90% reduction, p<0.0001). The α_4_ subunit, which forms heterodimers with integrin β_1_ as well as β_7_, was reduced ∼ 30% in SHP-1^−/−^ B cells (p<0.0001) reflecting the reduction in α_4_β_7_. In contrast, we found no reduction for integrins α_L_, β_1,_ and β_2_ (**Figure 1A**). We observed a similar and selective decrease of integrin β_7_ in CD4^+^ and CD8^+^ T cells (**Figure 1A**). Full SHP-1 deficiency leading to profound changes in the phenotype and homeostasis of mature B cells(Pao et al., 2007) and naïve T cells (Martinez et al., 2016; Mercadante and Lorenz, 2017), we repeated the above experiments with heterozygous viable motheaten mice (+/me^V^, or SHP-1^+/−^). Despite some residual SHP-1 activity, SHP1^+/−^ B and T cells displayed a selective reduction of α_4_, β_7_, and α_4_β_7_ integrins while expressing normal levels of α_L_, β_1_, and β_2_ (**Figure 1B**). These data suggest that SHP-1 positively regulates the cell surface levels of integrin α_4_β_7_ on lymphocytes without affecting other integrins.

**Figure 1.**
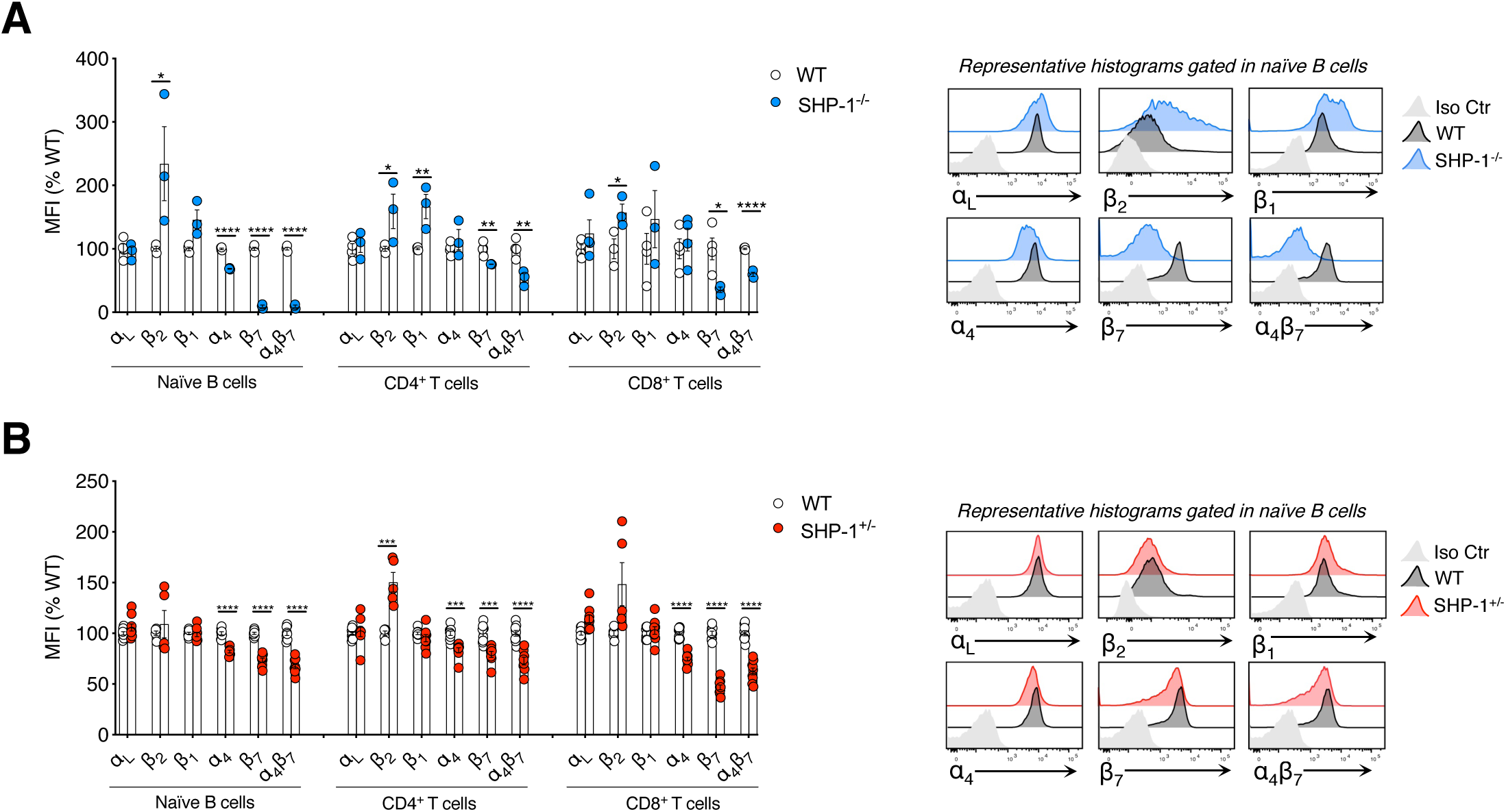
Selective reduction of integrin α_4_β_7_ on SHP-1 deficient B and T cells **(A)** Flow cytometry of WT or SHP-1^−/−^ naïve B cells (left panel), CD4^+^ T cells (mid-panel), and CD8^+^ T cells (right panel) isolated from spleens and stained for α_L_, β_2_, β_1_, α_4_, β_7_, or α_4_β_7_. Shown are pooled data (mean ± SEM) from n=3 experiments with 3-4 animals per group total. For each animal within one experiment, the MFI of the integrin staining was expressed as a percentage of the mean MFI of the WT group. Representative histogram overlays are shown (Iso Ctr: Isotype control). **(B)** Flow cytometry of WT or SHP-1^+/−^ naïve B cells (left panel), CD4^+^ T cells (mid-panel), and CD8^+^ T cells (right panel) isolated from spleens and stained for α_L_, β_2,_ β_1_, α_4_, β_7_, or α_4_β_7_. Shown are pooled data (mean ± SEM) from n=4 experiments with 4-9 animals per group total. Groups were compared by Student’s t-test with *P<0.05; **P< 0.01; ***P<0.001; ****P<0.0001.

### CD22 mediates the SHP-1-dependent α_4_β_7_ augmentation in B cells

The selective effect of SHP-1 haploinsufficiency on B cells suggested potential involvement of CD22, a B cell specific Siglec known to recruit SHP-1 to the plasma membrane through interactions with its three cytoplasmic ITIM sequences (Blasioli et al., 1999; Campbell and Klinman, 1995; Law et al., 1996). B cells from CD22^−/−^ mice showed reduced α_4_, β_7_, and α_4_β_7_ expression, but normal expression of α_L_, or β_1,_ (**Figure 2B**). B cells isolated from the bone marrow (BM), peripheral lymph nodes (PLNs) and Peyer’s patches (PPs) of CD22^−/−^ animals displayed a similar reduction in α_4_β_7_ (**Figure S1A**). Consistent with selective expression of CD22 by B cells, CD22 deficiency had no effect on CD4^+^ T cell integrins (**Figure S2**). mRNA expression of the α_4_ and β_7_ subunits in wild-type and CD22^−/−^ B cells was unchanged, ruling out an effect of CD22 deficiency on ITGA4 and ITGB7 gene expression (**Figure S1B**). Together, the data suggest that CD22 mediates the SHP-1 effect in B cells.

**Figure 2.**
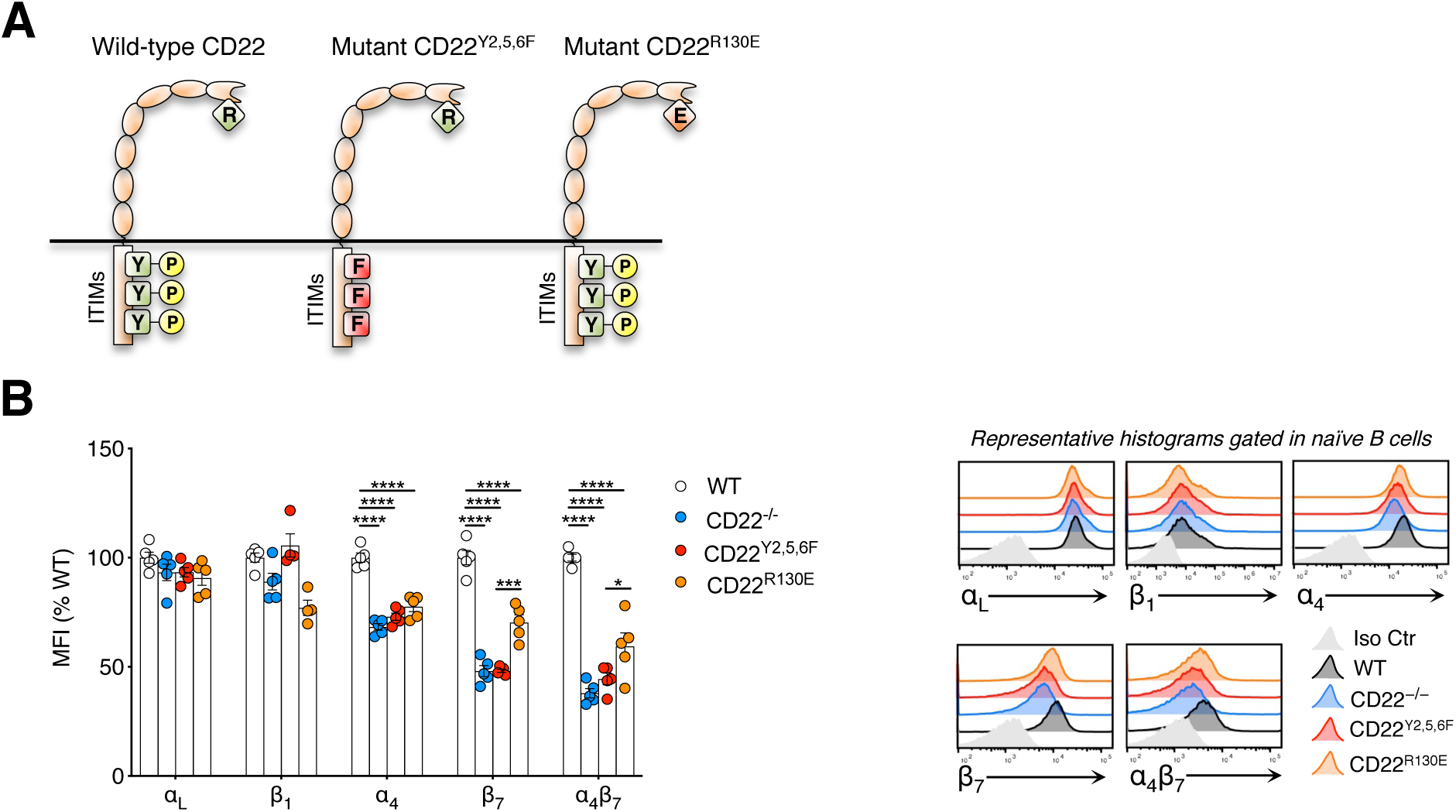
CD22 mediates the SHP-1-dependent α_4_β_7_ augmentation in B cells **(A)** Scheme of wild-type CD22, CD22^Y2,5,6F^, and CD22^R130E^ mutants. CD22^Y2,5,6F^ mutant displays a normal lectin binding but key tyrosine residues (Y) of the three ITIM domains were replaced by phenylalanine (F) to impair SHP-1 recruitment to the phosporylated ITIMs. CD22^R130E^ mutant displays normal ITIMs but one arginine (R) residue was replaced by a glutamic acid (E) to impair the lectin binding activity of CD22. **(B)** Flow cytometry of WT and CD22 mutant B cell isolated from spleens and stained for α_L_, β_1_, α_4_, β_7_, or α_4_β_7_. Shown are pooled data (mean ± SEM) from n=3 experiments with 7 animals per group total analyzed and presented as in Figure 1. Groups were compared by one way ANOVA with Dunnett’s or Tukey’s multiple comparison test. *P<0.05; ***P<0.001 and ****P<0.0001

To ask whether targeting SHP-1 to the CD22 ITIMs is required for the regulation of α_4_β_7_ levels, we used transgenic animals (CD22^Y2,5,6F^) in which the CD22 ITIM signaling domains has been mutated (Müller et al., 2013) to prevent SHP-1 binding and downstream CD22 signaling (**Figure 2A**). These mutants express normal levels of CD22 (Müller et al., 2013). CD22 ^Y2,5,6F^ B cells displayed levels of α_4_, β_7_, and α_4_β_7_ as severely reduced as on CD22^−/−^ B cells used here as controls, again with no effect on the other integrin levels (**Figure 2B**). Thus, the recruitment of SHP-1 to the CD22 ITIMs is necessary to the effects.

CD22 is a lectin that interacts in *cis* with α2-6 sialic acid-decorated glycoproteins, affecting its distribution and motility on the B cell surface. To assess a potential role for the CD22 lectin-carbohydrate interactions in cell autonomous α_4_β_7_ regulation, we assessed integrin expression by B cells expressing a mutated lectin domain (CD22^R130E^) that prevents α2-6 sialic acid binding (Müller et al., 2013) (**Figure 2A**). CD22^R130E^ B cells expressed CD22 at wild-type levels, and displayed a significant reduction in α_4_β_7_ levels (**Figure 2B**), although the effect was less dramatic than that of CD22 deficiency or ITIM mutations. Intermediate reduction in α_4_β_7_ was also observed in B cells from St6Gal1^−/−^ mice, which lack α2-6 sialyltransferase activity (Stamenkovic et al., 1991, 1992) and thus lack cognate CD22 α2-6-modified ligands (**Figure S3**). The result suggests that CD22-carbohydrates interactions at the cell surface contribute to normal α_4_β_7_ expression, potentially by controlling domain specific SHP-1 activity.

### CD22-deficiency increases β_7_ endocytosis in B cells via SHP-1 and ligand-recognition

Cell surface receptor expression can be regulated by endocytosis and recycling (Cullen and Steinberg, 2018; Grant and Donaldson, 2009). We asked if SHP-1 or CD22 might regulate β_7_ endocytosis. We assessed receptor internalization by flow cytometry using pHrodo-Red, a dye whose fluorescence increases within the acidic pH of endocytic vesicles. We compared the effects of CD22/SHP-1 deficiency on endocytosis of β_7_, of β_1_ integrin which is not affected by CD22 or SHP-1 (**Figure 1B** and **2A**), and transferrin (Tf) included as an irrelevant control that undergoes endocytic recycling. We incubated splenocytes for one hour at 37°C to enable endocytosis, in the presence of pHrodo-red-sconjugated Tf, or anti-β_7,_ anti-β_1,_ or isotype matched control Abs. Background was defined by staining of cells with the pHrodo-Red constructs incubated at 4°C to prevent endocytosis. Endocytosis, defined by the internalization induced (background corrected) pHrodo-Red signal, was normalized to cell surface expression of each antigen at 4°C to yield a Relative Endocytosis Ratio (RER, **Figure 3A**).

**Figure 3.**
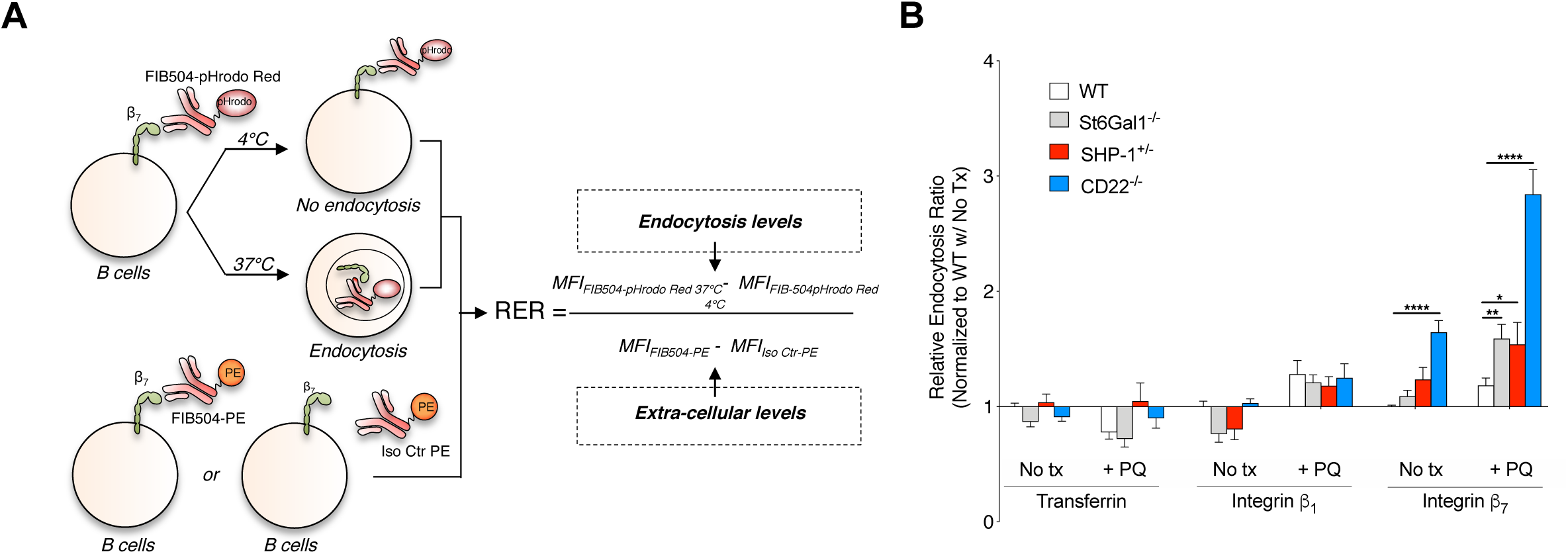
CD22-deficiency increases β_7_ endocytosis in B cells via SHP-1 and ligand-recognition **(A)** Splenocytes staining with anti-CD19 and anti-IgD (to identify B cells) and pHrodo-Red-conjugated tranferrin (Tf), HMβ1-1 (anti-β_1_), or FIB504 (anti-β_7_) antibodies at either 4°C (no endocytosis) or 37°C (endocytosis) was used to calculate endocytosis levels (i.e. MFI pHrodo-Red staining at 37°C – MFI pHrodo-Red staining at 4°C). Staining at 4°C with Phycoerythrin (PE)-conjugated RI7217 (anti-Transferrin Receptor 1, TfR1), HMβ1-1, FIB504 or the matching PE-conjugated isotype control antibodies was used to calculate levels of extra-cellular TfR1, β_1_, and β_7_ levels (i.e. MFI PE staining at 4°C – MFI Isotype Control staining at 4°C). For each molecule (illustrated with β_7_ in panel (A)) and each experiment, the Relative Endocytosis Ratio (RER) was calculated by normalizing endocytosis levels to extra-cellular levels. **(B)** RER of Tf, β_1_, and β_7_ in WT, CD22^−/−^, SHP-1^+/−^, and St6Gal1^−/−^ B cells with (100µM) or without primaquine (PQ) treatment. The mean RER of the WT B cells group without PQ (No Tx) was set to 1, and all data were normalized to that mean value. Shown are pooled data (mean ± SEM) from n=3 independent experiments with 6 animals per group total. *P<0.01, **P<0.01; and ****P<0.0001 (Two-way ANOVA with Sidak’s multiple comparison test).

CD22-deficient B cells displayed a significantly higher RER for integrin β_7_ compared to wild-type B cells (1.65 fold higher, p<0.0001). CD22-deficiency did not alter the basal endocytosis of Tf, or of integrin β_1_ used as negative controls (**Figure 3B**), while we observed no endocytosis of the isotype control Ab (data not shown). SHP-1 haploinsufficient B cells showed a similar shift than CD22 knock-out, but not reaching significance. Since recycling of receptors back to the cell surface could reduce measured internalization, we also studied endocytosis in presence of the anti-malarial drug primaquine (PQ) which inhibits the recycling of endocytosed proteins to the plasma membrane(Van Weert et al., 2000). Treatment of B cells with PQ significantly increased measured internalization. β_7_ internalization was significantly increased in SHP-1^+/−^ B cells (RER 1.54 vs. 1.18, p<0.05) and in St6gal1^−/−^ B cells (1.59 vs. 1.18, p<0.01); and PQ treatment significantly enhanced the difference between CD22^−/−^ and wild-type B cells (2.84 vs. 1.18, p<0.0001) (**Figure 3B**). Thus CD22 and SHP-1 similarly and selectively inhibit β_7_ endocytosis in B cells.

### Functional consequences of SHP-1-CD22 dependent β_7_ augmentation

The integrin α_4_β_7_ plays a key role in cell-cell and cell-substrate adhesive interactions involved in gastrointestinal immune responses, including lymphocytes interactions with matrix components and stromal cells and lymphocyte homing to the intestines. To assess the consequence of SHP1- and CD22-dependent α_4_β_7_ regulation in functional assays, we initially assessed the short term homing of B cells into Peyer’s patches and mesenteric lymph nodes (MLN) which express the dedicated α_4_β_7_ ligand MAdCAM-1. Recipient wild-type mice received splenocytes from wild-type and mutant donors labeled with different cell tracker dyes and mixed at a 1:1:1 ratio as confirmed by flow cytometry. Two hours later, we enumerated cells localized to PP and MLN by flow cytometry (**Figure 4A**). In agreement with the reduction of α_4_β_7_, SHP-1^+/−^ B cells, CD22^−/−^ B cells, CD22 ^Y2,5,6F^ ITIM mutant B cells, and CD22^R130E^ lectin mutant B cells all homed poorly to PP, displaying a similar ∼60% reduction in recruitment compared to WT B cells (**Figure 4B and 4C**). We observed an intermediate defect for all mutant B cells in homing to the MLN that is consistent with MAdCAM-1 expression while homing to PLN, which is independent of α_4_β_7_, was normal (**Figure 4B and 4C**). CD22 deficiency did not affect CD4^+^ T cell homing to PP while SHP-1 haplodeficiency compromised CD4^+^ T cell homing (**Figure S4**) consistent with reduced β_7_ levels (**Figure 1B)**. SHP-1^+/−^ CD4^+^ T cells also poorly localized to PLN (**Figure S4**). This may reflect the known role of SHP-1 in modulating β_2_ integrin expression in T cells (Sauer et al., 2016).

**Figure 4.**
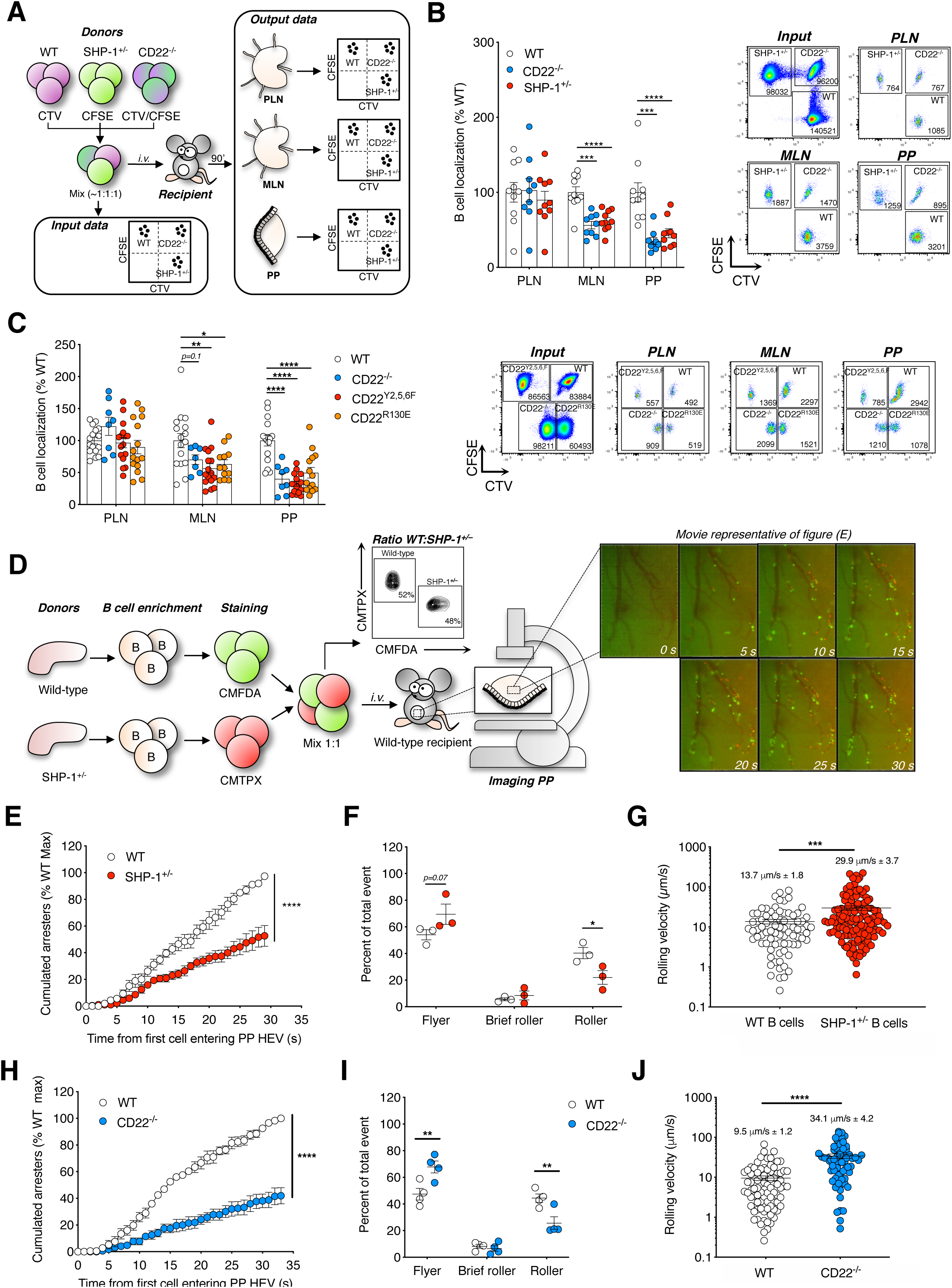
Functional assays reveals defective homing, rolling, and arrest of SHP-1^+/−^ B cells and CD22 mutant B cells to PP **(A)** WT, CD22^−/−^, or SHP-1^+/–^ splenocytes labeled with CFSE, or CellTracker Violet (CTV) or both were injected i.v. into a recipient WT mouse. PLN, MLN or PP cells isolated from the recipient were stained with anti-CD19 and anti-IgD for quantification of short-term (90 min) homing of B cells. For each donor and each organ, the number of isolated B cells (Output) was normalized to the number of injected B cells (Input) to yield a B cell localization ratio. **(B-C)** Localization of WT, CD22^−/−^ and SHP-1^+/–^ B cells (B) or WT, CD22^−/−^, CD22^Y2,5,6F^, and CD22^R130E^ (C) in PLN, MLN and PP after homing assays. Data are shown as a percentage of the mean localization ratio of the WT group. Shown are pooled data (mean ± SEM) of n=3-5 experiments with 11-16 mice per group total. Representative dot plots gated on live B cells are shown **(D)** B cells purified from WT and either SHP-1^+/–^ or CD22^−/−^ splenocytes were stained with cell trackers, mixed at a 1:1 ratio, and injected i.v. into a wild-type recipient. The interactions between B cells and PP-HEVs were visualized *in situ* on one isolated PP. **(E-J)** Data in panels (E,F,H,I) represent the mean ± SEM of three independent experiments. **(E,H)** Number of WT vs. SHP-1^+/−^ (E) or WT vs. CD22^−/−^ (H) arresters on PP HEVs shown second per second from the first cell entering the HEVs. The total amount of WT B cells arresters at the end each experiment (i.e. WT B cells max; ∼ 40–80 cells in average) was set to 100, and the data expressed as a percentage of this total. Representative images of one movie are shown in panel (D) with WT in green and SHP-1^+/−^ in red **(F,I)** The behavior (i.e. flyer, brief roller, or roller as defined in **Figure S5**) of each cell entering the HEVs was analysed in 3–4 representative PP-HEVs per experiment. Results are shown as a percentage of the total numbers of cells entering HEVs (∼ 250–300 total cells analyzed per group) **(G,J)** Average rolling velocity of representative rollers. Groups were compared by one way ANOVA (B,C), paired one-tailed Student’s t-test (E,H), and unpaired one-tailed Student’s t-test (F,G,I,J). \**P* < 0.05, **P<0.01, \*\*\**P* < 0.001, \*\*\*\**P* < 0.0001.

α_4_β_7_ mediates both activation-independent tethering and rolling on PP, and chemokine/integrin activation-dependent arrest in combination with LFA-1 (Bargatze et al., 1995). To assess the effects of the CD22-SHP-1 axis on these steps, we visualized B and T cell behavior in PP by intravital microscopy. We purified B or T cells from wild-type, SHP-1^+/−^ and CD22^−/−^ donors, labeled them with different cell tracker dyes, mixed mutant and WT cells at a 1:1 ratio as confirmed by flow cytometry, and transferred the cells into a recipient wild-type mouse while imaging the PP (**Figure 4D**). We captured high frame rate movies (40 frames per second) to study the behavior of all cells entering HEV within the field of view during the recording (i.e. ∼30-40 s). We stratified the cells into four groups based on their interactions with HEV: freely flowing cells that failed to interact detectably, termed flyers, appeared as streaks due to their high velocity during the time of exposure, and passed through the HEVs at a mean velocity of ∼ 1000 µm/s (**Figure S5A** and **Supplemental Video 1**). Cell capture on the vessel wall could be visualized as soon as the cell looks round and bright as a result of reduced velocity. Cells attaching briefly (< 1s) before detaching and flying through the HEV were described as brief rollers (**Figure S5B and S5C** and **Supplemental Video 2**). Cells interacting for more than 1 sec were called rollers (**Figure S5B and S5C** and **Supplemental Video 3**). Among rollers, cells static for more than 2 s at the end of the recording were considered arresters.

At the end of ∼30 seconds of observation, the number of SHP-1**^+/–^** and CD22^−/−^ B cell arresters was ∼50% and ∼60% lower (p<0.0001) respectively than wild-type B cells (**Figure 4E and 4H; Supplemental Video 4 and 5**). The reduced frequency of arrest correlated with inefficient initial capture or tethering, as well as faster rolling (looser interaction) of SHP-1**^+/–^** and CD22^−/−^ B cells. We observed a reduced frequency of rolling by SHP-1**^+/–^** and CD22^−/−^ B cells as compared to wild-type control cells with a corresponding increase in non-interacting flyers (**Figure 4F and 4I**). Among rollers, the average rolling velocity of SHP-1**^+/–^** and CD22^−/−^ B cells was ∼2-3 times higher than wild-type control (**Figure 4G and 4J; Supplemental Video 6 and 7**). The total number of mutant cells and wild-type B cells entering PP-HEVs (including roller, brief rollers, or flyers) was similar in each experiment, thus ruling out distortion of results by differences in retention of mutant cells in other organs (**Figure S5D**). Also the effects of CD22 deficiency and SHP-1 haploinsufficiency on tethering and rolling velocity are mimicked by experimental reduction in surface α_4_β_7_ available for interactions (**Figure S6**)

These data suggest that SHP-1-driven CD22 regulation of β_7_ in B cells augments tethering and slow rolling, consistent with a primary role in controlling cell surface expression levels of integrin α_4_β_7_.

### Defects in intestinal vs. systemic antibody responses in CD22-deficient mice

In addition to its roles in gut lymphocyte homing, α_4_β_7_ mediates cell-cell interactions of lymphocytes with MAdCAM-1 expressed by follicular dendritic cells in GALT (Butcher et al., 1997). It also supports interactions of α_4_β_7_-expressing lymphocytes with other ligands including α_4_ on adjacent immune cells, fibronectin in the extracellular matrix, and VCAM-1 (Altevogt et al., 1995; Berlin et al., 1993). Since α_4_β_7_ is preferentially expressed by memory and effector B cells involved in gut responses (Williams et al., 1998), we reasoned that CD22-dependent α_4_β_7_ expression could have a selective role in intestinal vs. systemic antibody responses.

To test this hypothesis, we immunized cohorts of wild-type and CD22-deficient animals with a single dose of a potent immunogen, Cholera Toxin B (CTB), comparing the local and systemic humoral responses after oral, intra-nasal, and intramuscular immunizations. Two weeks after CTB administration, we measured systemic Ag-specific IgA and IgG in the serum. We observed a significant reduction of CTB specific IgA and IgG levels in CD22-deficient animals immunized orally (**Figure 5A** and **Figure 5B**). Importantly, total IgA and IgG levels in the serum after oral, nasal, and intramuscular immunizations were not different in CD22 deficient vs. WT animals ruling out a global inhibition of B cell responses (**Figure S7**).

**Figure 5.**
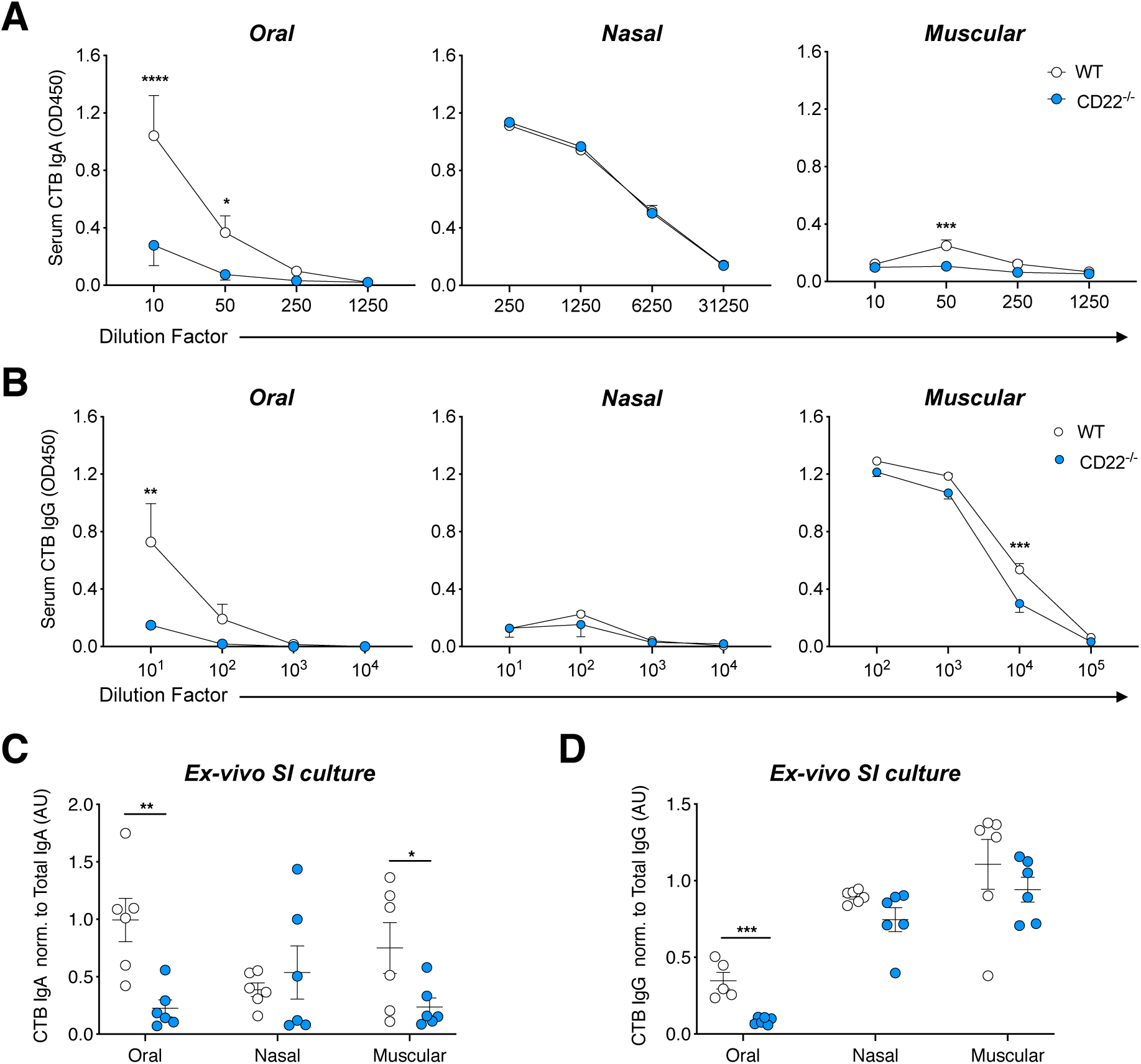
Defects in intestinal responses to oral antigen in CD22-deficient mice. Cohorts of WT or CD22^−/−^ mice were immunized with Cholera Toxin B (CTB) via the Oral, intra-nasal (Nasal), or intra-muscular (Muscular) routes for two weeks. **(A-B)** The serum levels of CTB-specific IgA **(A)** or IgG **(B)** were measured by ELISA and expressed as net OD450. Shown are the mean ± SEM with n=3 mice per group from one out of two independent experiments with similar results. *P<0.05, ***P<0.001 and ****P<0.0001 (Two-way ANOVA with Sidak’s multiple comparison test). **(C-D)** SI segments were cultured *ex-vivo* for three days to titer the quantity of secreted CTB-specific IgA **(C)** and IgG **(D)** produced by ELISA. For each sample, the titer (OD450) of CTB-specific Ig response was normalized to the titer (OD450) of the total Ig response. Shown are pooled data (mean ± SEM) from n=2 experiments with 6 mice per group total. *P<0.05 and ***P<0.001 (Student’s t-test).

To test whether the systemic reduction of CTB IgA and IgG levels after oral immunization in CD22^−/−^ mice correlate with a deficit in local production in the gut, we analyzed the CTB-specific IgA and IgG levels in *ex-vivo* culture of SI segments. Interestingly, we found a strong decrease of CTB-specific IgA (**Figure 5C**) and IgG levels (**Figure 5D**) in CD22^−/−^ SI segments after oral immunization while a moderate effect after intra-nasal or intra-muscular immunizations (**Figure 5C and 5D**). Together, these data confirm a selective role for CD22 in the intestinal immune response, consistent with its role in regulating the gut lymphocyte integrin α_4_β_7_.

### CD22 deficiency delays the protective immune response to rotavirus infection

We further investigated the specific role for CD22 in the intestinal response in a mouse model of rotavirus (RV) infection. RV selectively infects intestinal epithelial cells, leading to strong immune responses in GALT; and the local production of RV-specific IgA is critical for protection (Burns et al., 1996; Vo et al., 2011). Young pups, more susceptible to infection than adults, were infected with RV. We measured total and RV-specific IgA and IgG in the feces daily. Ten days post infection (p.i.), wild-type animals started to mount a robust RV-specific intestinal IgA and IgG responses (**Figure 6A** and **Figure S8A**), which correlated with a significant reduction of the fecal viral shedding by day 12 p.i. and a full resolution of RV infection by day 15 p.i. (**Figure 6B** and **Figure S8B**). In contrast, CD22-deficient animals had a dramatically reduced RV-specific IgA response in the feces (**Figure 6A**), and a delay in the resolution as indicated by residual viral shedding at day 12 p.i. (**Figure 6B)**. Virus was eliminated by day 15 p.i. in both WT and CD22^−/−^ mice (**Figure S8B**).

**Figure 6.**
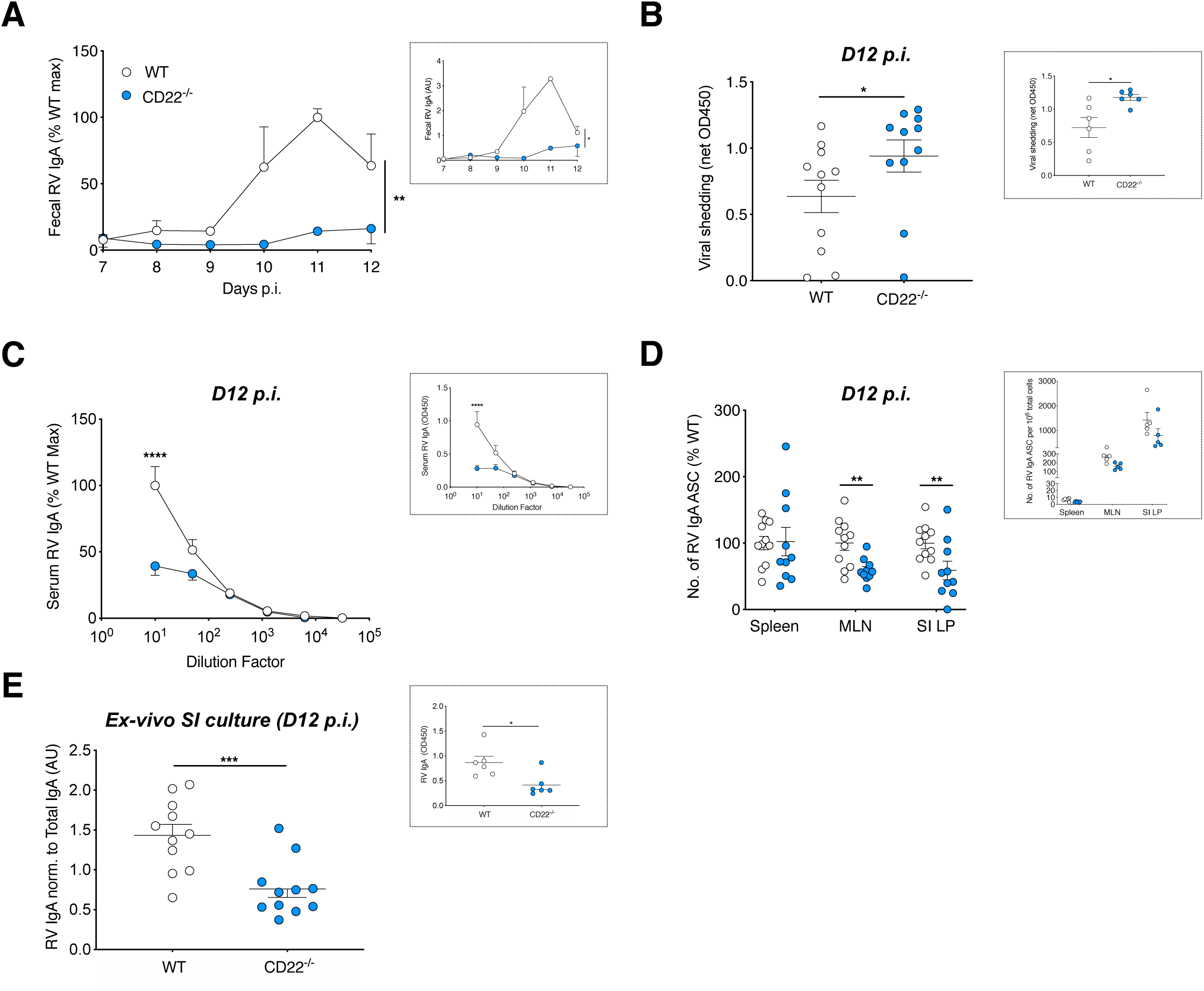
Delayed protective immune response to RV infection in CD22-deficient animals. **(A)** Five days old WT or CD22^−/−^ pups were orally gavaged with the RV strain EW. The production of RV-specific IgA and total IgA in fecal samples was measured by ELISA up to twelve days post-infection (p.i.). For each individual animal within one experiment, the ratio of RV IgA to total IgA levels was expressed as a percentage of the maximal ratio measured in the WT group. Shown are pooled data (mean ± SEM) from n=3 independent experiments with 5-10 animals per group total. Boxed inset: ratios of RV IgA to total IgA shown as arbitrary unit (AU) in one representative experiment. **(B-E)** In separate experiments, five days old WT and CD22^−/−^ pups were orally inoculated with RV EW and sacrificed at day 12 p.i. Shown in panels (B-E) are pooled data (mean ± SEM) from n=2 independent experiments with 10-11 mice per group. Boxed insets show the result of one out of the two experiments **(B)** Fecal RV Ag shedding as measured by ELISA and expressed as net OD450, normalized to the sample weight. **(C)** The serum levels of RV-specific IgA were measured by ELISA and expressed as net OD450. For each individual animal within one experiment, the net OD450 was expressed as a percentage of the maximal net OD450 measured in the WT group at the lowest serum dilution (1:10). **(D)** The numbers of RV-specific IgA antibody secreting B cells (ASC) were measured in the spleen, mesenteric lymph nodes (MLN), and small intestine lamina propria (SI LP) by ELISPOT. Shown are numbers of RV IgA ASC per 10^6^ total cells expressed as a percentage of the WT group mean. **(E)** Ratio of RV IgA to total IgA in *ex-vivo* SI segments cultures (AU). Groups were compared by paired Student’s t-test (A), unpaired one-tailed Student’s t-test (B,D,E) and Two-way ANOVA with Sidak’s multiple comparison test (C). \**P* < 0.05, **P<0.01, \*\*\**P* < 0.001, \*\*\*\**P* < 0.0001.

At the peak of the fecal Ab response (day 12 p.i.), we observed reduced RV-specific IgA titers in the serum of CD22-deficient animals (**Figure 6C**) while RV-specific IgG, total IgG and total IgA levels were similar to wild-type (**Figure S8C, S8E, and S8F**). CD22^−/−^ animals also had fewer RV-specific IgA antibody secreting B cells in the mesenteric lymph nodes (MLN) and the small intestine lamina propria (SI LP) than WT mice at day 12 p.i., while there were no differences in the spleen (**Figure 6D**). To determine whether the reduction of RV-specific fecal antibody in CD22^−/−^ mice reflected a deficit in local production in the gut, we analyzed RV-specific IgA and IgG levels in *ex-vivo* culture of SI segments. We found a two-fold decrease in the RV-specific IgA and IgG levels in CD22-deficient animals as compared to wild-type controls **(Figure 6E** and **Figure S8D**), which correlates with higher viral shedding in the feces at day 12 p.i. as mentioned above (**Figure 6B**). Together, these data highlight a significant role of B cell CD22 in the mucosal immune response to enteric pathogens.

## Discussion

In this study, we report the cell autonomous regulation of the gut homing receptor α_4_β_7_ by SHP-1 on B and T lymphocytes. The effect is specific to the intestinal lymphocyte integrin α_4_β_7_: SHP-1 deficiency reduces β_7_ but not β_2_ or β_1_ integrin expression with a dramatic effect in B cells. We show that SHP-1 inhibits α_4_β_7_ endocytosis thus maintaining cell surface α_4_β_7_ expression. We also show that SHP-1 is required for normal α_4_β_7_-dependent tethering and arrest of B cells to PP-HEVs. The B cell specific lectin CD22 mediates the dramatic SHP-1 dependent β_7_ regulation seen on B lymphocytes. Both the carbohydrate-binding lectin activity and cytoplasmic SHP-1-binding ITIM domains of CD22 are important for surface α_4_β_7_ expression. We suggest a model in which the CD22 ITIM domains can recruit and target the SHP-1 phosphatase to the cell surface to regulate β_7_ endocytosis. We also demonstrate a selective role for CD22 in intestinal antibody responses and mucosal immunity in multiple models. The results define a novel mechanism for differential control of integrin function and an unexpected and selective role for CD22 in intestinal immunity.

We showed that the tyrosine phosphatase SHP-1 specifically controls β_7_ levels at the cell surface: haploinsufficiency of SHP-1 enhances β_7_ but not β_1_ endocytosis, leading to reduced surface α_4_β_7_ expression. SHP-1 is an essential immune regulator of signaling pathways including those of receptor tyrosine kinase, cytokine or chemokine receptors, and T and B cell receptors (BCR) (Neel et al., 2003; Zhang et al., 2000). Amongst its wide spectrum of activities, SHP-1 often negatively regulates its target to prevent overactivation of immune modalities.

SHP-1 regulates intracellular signaling, and prevents hyper activation of the β_2_ integrins in macrophages (Roach et al., 1998), neutrophils (Kruger et al., 2000) and dendritic cells (Abram et al., 2013). Βeta integrin dephosphorylation by tyrosine phosphatases in general is a well documented and highly conserved mechanism controlling integrin activation(Anthis et al., 2009; Legate and Fassler, 2008). However the role of tyrosine phosphatases in the control of integrin endocytosis and surface expression has not been reported. Importantly, SHP-1 controls β_7_ levels in both B and T cells. In B cells, CD22 recruits SHP-1 via its ITIM domains to the plasma membrane, effectively increasing its access to tyrosine phosphorylated membrane associated targets. This suggests that T cells like B cells could adapt T cell-expressed ITIM-based accessory proteins for the specific SHP-1-dependent enhancement of β_7_ levels. In contrast to the reduction in β_7_, β_1_ and β_2_ integrins are actually more highly expressed at the cell surface of SHP-1^−/−^ B and T cells than on wild-type cells, confirming the integrin selectivity of its actions and suggesting further complexity in its potential roles in differential lymphocyte adhesion and homing.

Mechanistically, we found that, in addition to the ITIM docking motifs in the cytoplasmic tail, the lectin-carbohydrate binding activity of CD22 also contributes to α_4_β_7_ regulation, although the reduction of α_4_β_7_ levels on B cells, and the inhibition of PP homing were less dramatic in the lectin mutant than with ITIM deficiency. On B cells, CD22 associates in *cis* with α2-6 Sia modified membrane proteins including CD22 itself and CD45: these *cis*-interactions can bring ITIM-bound SHP-1 phosphatase in close proximity to target substrates, altering the phosphorylation state and function of associated molecules (Enterina et al., 2019; Meyer et al., 2018). Using a biotin-tyramide-based proximity labeling method (Alborzian Deh Sheikh et al., 2018) to label molecules in close proximity to CD22, we reproduced the finding that CD45 and CD22 are α2-6 Sia-dependent CD22 cis-ligands, but we did not detect interaction with α_4_β_7_ (data not shown), suggesting either that, such interactions are transient (Collins et al., 2004) or that CD22 acts through intermediate signaling molecules to regulate α_4_β_7_. Lectin interactions could also contribute to the domain localization of CD22 on the cell surface. In CD22^R130E^ B cells, CD22 itself is known to be more phosphorylated, recruits more SHP-1, and has higher lateral mobility than in wild-type B cells (Gasparrini et al., 2015; Müller et al., 2013). This enhanced mobility could counter the loss of any targeting function of *cis* lectin-CHO interactions, thus explaining the limited effect of lectin vs. ITIM mutations. Taken together, the data suggest a model in which SHP-1 is targeted to the cell surface by the CD22 ITIM domains to inhibit endocytosis of the integrin β_7,_ leading to enhanced α_4_β_7_ surface expression and increased intestinal homing of B cells (**Figure S10**). How CD22 and SHP-1 select β_7_ vs. β_1_ or β_2_ for endocytosis remains to be determined. The SHP-1 phosphatase may selectively recognize phosphotyrosine motifs of the β_7_ chain cytoplasmic tail that bind endocytosis adaptor proteins. Alternatively (or in addition), selective targeting or association of CD22-SHP1 to α_4_β_7_ remains a possibility. Further studies may shed light on the exact mechanism of CD22/SHP-1 specificity for β_7_.

This selective regulation of the intestinal homing receptor α_4_β_7_ by CD22 and SHP-1 has significant implications for intestinal immune responses. B cell activation in the GALT drives production of local and systemic antibodies to intestinal antigens and pathogens. We evaluated the antibody response in CD22 deficient mice that show grossly normal B cell development (Nitschke et al., 1997). We observed a profound defect in the intestinal and systemic Ag-specific IgA and IgG levels to oral immunizations with Cholera-Toxin B in CD22 deficient animals, whereas the IgG response to nasal or muscular immunizations was preserved. The results closely recapitulate previous findings with β_7_-deficiency or α_4_β_7_ blockade: Shippers and colleagues reported a similar reduction of intestinal Ag-specific IgA levels in β_7_-deficient mice immunized orally with cholera toxin (Schippers et al., 2012). In humans, an antibody blocking the α_4_β_7_ heterodimer also impaired the Ab responses to an oral Ag while not changing the response to a systemic immunization (Schippers et al., 2012). The poor mucosal immune response in CD22 deficient mice is likely the result of the defective α_4_β_7_-dependent homing of B cells to the GALT and α_4_β_7_-dependent B cells interactions in the inductive or effector sites of the gut. We also observed a variable reduction of Ag-specific IgA, but not IgG levels in the GALT after systemic immunization. Shippers et al. also observed a moderate reduction of intestinal Ag-specific IgA levels, but not IgG, in β_7_-deficient animals after systemic immunization with NP-CG (Schippers et al., 2012). This could reflect a contribution of α_4_β_7_ to (infrequent) IgA isotype switching in systemic lymphoid tissues. More likely is generation of IgA ASC in GALT: after cutaneous immunization, antigen presenting cells can migrate to GALT including PP, where they can induce mucosal immunity (Enioutina et al., 2008).

Intestinal antibody responses mediate protection against intestinal pathogens including rotavirus, a reovirus that is the most common cause of childhood diarrheal illness worldwide. Rotavirus selectively infects small intestinal epithelial cells, providing a model for the gut specific immune response (Feng et al., 1997). Consistent with its importance to α_4_β_7_ expression and intestinal IgA production, CD22 plays an important role in the rotavirus response: CD22 deficiency caused a significant reduction in viral clearance, comparable to that reported in mice with a complete lack of B cells (Marcelin et al., 2011; McNeal et al., 1995), or lacking IgA (Blutt et al., 2012), or lacking GALT (Lopatin et al., 2012) itself. Given the diversity of innate and adaptive mechanisms that can affect viral clearance (Feng et al., 2008; Franco and Greenberg, 1995a), the effect of CD22 deficiency is quite significant and likely reflects global influences of CD22-dependent α_4_β_7_ functions in the intestinal immune response.

The specific targeting of immune cells to the GALT involves distinct molecular signatures on HEVs and venules as well as lymphocytes. α_4_β_7_ mediates B cell recognition of GALT HEV expressed MAdCAM-1. We previously showed that PP HEVs, but not PLN HEVs, also express high levels of *St6Gal1*, generating functional ligands for the B cell lectin CD22. St6Gal1^−/−^ PP-HEV are defective at recruiting CD22-expressing B cells in short-term homing assays (Lee et al., 2014), likely as a result of trans-interactions between CD22 and α2-6 Sia decorated glycans on HEVs. Yet we showed here that even in absence of CD22-ligand on PP-HEVs, reduced α_4_β_7_ on CD22^−/−^ B cell significantly impacts homing to the PP (**Figure S9**). Intravital microscopy in ligand-deficient mice revealed that wild-type B cell tethering and rolling velocity on St6Gal1^−/−^ PP-HEV are normal: only definitive arrest is compromised (unpublished observations). We hypothesize that CD22 ligation to α2-6 Sia-modified MAdCAM-1 may create a synapse between CD22, MAdCAM-1 and α_4_β_7_ facilitating chemokine-driven activation of α_4_β_7_ and arrest of the B cell. An analogous mechanism controls the arrest of neutrophils: heterophilic ligation of endothelial cell expressed-CD99 to paired immunoglobulin-like receptors (PILRs) enhances the chemokine-induced β_2_-dependent arrest of neutrophils on ICAM-1 (Goswami et al., 2017). Future studies will likely shed the light on this complex mechanism. It is fascinating that the same molecule (namely CD22) has evolved into two separate functions (i) selective regulation of β_7_ expression and (ii) trans-ligation to PP-HEVs heterophilic ligands to serve the same goal, B cell homing to the GALT. This dual function, besides its role in B cell tolerance, makes CD22 a unique example of a Swiss army knife molecule for leukocyte homing.

In conclusion, our study reveals a novel role for SHP-1 in selective regulation of integrin endocytosis, and unravels a CD22-dependent SHP-1-driven mechanism of α_4_β_7_ regulation in B cells and a BCR-independent role for CD22 in enhancing intestinal immune responses.

## Author contribution

R.B. conceptualized the study, designed, performed, analyzed the majority of the experiments, and wrote the manuscript. C.B. performed experiments involving the CD22^Y256F^ and CD22^R130E^ transgenic animals; N.F. performed oral RV infections, and helped conceptualizing and designing the RV studies; J.B. and J.C. contributed to the analysis of the video-microcopy experiments; B.O. performed gut preparations in the RV studies and helped with SI fragment cultures; A.M. shared his intravital microscopy expertise with R.B.; A.A.D.S. performed biotin-tyramide-based proximity labeling assays; C.A.L and C.L.A. provided the *motheathen* mice; T.T. contributed to conceptualizing and designing the proximity labeling assays; H.B.G. contributed to conceptualizing and designing the RV studies; M.S.M., K.L. and L.N. contributed to conceptualizing the study and provided intellectual input; E.C.B. guided, conceptualized and supervised the study, and wrote the manuscript.

## Acknowledgements

We thank J. Paulson for the CD22^−/−^ and St6Gal1^−/−^ mice; the members of the Butcher lab for discussions; H. Hadeiba for helpful discussions and for help with some experiments not included in this manuscript; M. Brennan for help with experiments; C. Garzon-Coral for designing and making re-usable dishes used for positioning animals in intravital imaging studies; J.L. Jang for the production of home-made Abs used in these studies; M. Bscheider for helping implementing the imaging softwares in the Butcher lab that were used to record and analyzed the video-microscopy; A. Scholz for helpful discussions; M. Lajevic for sharing protocols and expertise. This work was supported by NIH grants R37AI047822 and R01 AI130471 and award I01 BX-002919 from the Department of Veterans Affairs to E.C.B.; by the Swiss National Sciences Foundation grants P2GEP3_162055 and P300PA_174365 to R.B.; the DFG-funded TRR130 (project 04) to L.N.; JSPS Grant-in-Aid for Scientific Research 18H02610 and 19H04804 to T.T.; 1R01 AI125249 (NIH/NIAID), 1IO 1BX000158-01A1 (Veterans Affairs) to H.B.G.; and the Ramon Areces Foundation (Madrid, Spain) Postdoctoral Fellowship and Research Fellow Award (Crohn’s and Colitis Foundation of America) to B.O. M.S.M. acknowledges funding provided through NIAID (AI118842).

## Material and methods

### Reagents

Allophycocyanin-conjugated anti-mouse CD4 (RM4-5), phycoerythrin-indotricarbocyanine-conjugated anti-mouse CD3ε (145-2C11), peridinin-chlorophyll protein-Cyanine 5.5-conjugated anti-mouse IgD (11.26c.2a), phycoerythrin-conjugated anti-mouse CD19 (6D5), Brilliant violet 421-conjugated anti-mouse CD4 (RM4-5), phycoerythrin-conjugated armenian hamster IgG isotype control, phycoerythrin-conjugated anti-mouse CD49d (9C10), phycoerythrin-conjugated anti-mouse CD18 (M18/2), purified anti rat/mouse CD29 (HM β1-1) were from Biolegend. Fluorescein isothiocyanate-conjugated anti-mouse CD19 (1D3), fluorescein isothiocyanate-conjugated anti-mouse IgD (11-26), allophycocyanin-conjugated anti-mouse CD3e (145-2C11) was from eBioscience. Phycoerythrin-conjugated rat IgG2a,κ isotype control, phycoerythrin-conjugated anti-mouse CD11a (2D7), phycoerythrin-conjugated anti-mouse CD29 (HM β1-1), phycoerythrin-conjugated anti-mouse integrin β7 (M293), phycoerythrin-conjugated anti-mouse LPAM-1 (DATK32), purified rat anti-mouse CD16/CD32 (2.4G2), purified rat anti-mouse CD11a (2D7), allophycocyanin-cyanine 7 rat anti-mouse CD19 (1D3) were from BD. Purified anti-mouse/human integrin β7 (FIB504) was from BioXCell. Carboxyfluorescein succinimidyl ester (CFSE), CellTracker violet (CTV), CellTracker green (CMFDA), CellTracker red (CMTPX), Live/Dead Fixable Aqua dead cell staining kit, pHrodo™ iFL microscale protein labeling kits, High Capacity cDNA Reverse Transcription Kit, PowerSYBR**®** Green PCR Master Mix, CountBright absolute counting beads for flow cytometry were from Invitrogen. The mouse B cell isolation kit and mouse naïve CD4^+^ T cell isolation kit were from STEMCELL technologies. The purified rat anti-mouse α_4_β_7_ (DATK32) was in-house from hybridomas. The guinea pig anti-rotavirus hyperimmune serum, the rabbit anti-rotavirus hyperimmune serum, the rhesus rotavirus (RRV) stock, and purified double-layer RRV particles used in RV studies were gift from the Harry Greenberg’s lab. The horseradish peroxidase-conjugated goat anti-rabbit IgG, the peroxidase-conjugated anti-mouse IgA or IgG, and anti-mouse IgA, IgG, IgM polyconal antibodies were all from KPL. Cholera Toxin B, TMB, D-PBS were from Sigma. The RNeasy kit was from Qiagen. All reagents were titered and used according to the manufacturers’ recommendations.

### Mice

Mouse strains used: C57BL/6 wild-type mice, BALB/c wild-type mice, CD22^−/−^ mice(Nitschke et al., 1997), and St6Gal1^−/−^ mice(Hennet et al., 1998), were bred and housed in the animal facilities of the Veterans Affairs Palo Alto Health Care System, accredited by the Association for Assessment and Accreditation of Laboratory Animal Care. CD22^Y2,5,6F^ mice(Müller et al., 2013) and CD22^R130E^ mice(Müller et al., 2013) were bred and housed at the Department of Biology of the University of Erlangen; SHP-1^−/−^ and SHP-1^+/−^ mice(Coman and Bailey, 1984) were bred and housed at the University of California San Francisco. Unless otherwise stated, 6-12 weeks old mice were used in all experiments. All animal work was approved by the Institutional Animal Care and Use Committee at the Veterans Affairs Palo Alto Health Care System, or by relevant animal care committees at the University of Erlangen.

### Integrin levels by flow cytometry

Ten thousand splenocytes were stained with fluorescently labeled antibodies against CD4, CD3ε, CD19, IgD and either CD11a (α_L_), CD18 (β_2_) CD49d (α_4_), CD29 (β_1_), β7, LPAM-1 (α_4_β_7_), or the appropriate Isotype Control. The Fc receptors were blocked using Rat Serum and anti-CD16/32 prior to the staining. Dead cells were excluded by staining with 4’,6-diamidino-2-phénylindole (DAPI). For each sample, the median fluorescence intensity (MFI) of isotype control was substracted to the MFI of the integrin staining. The integrin staining MFI (background corrected) was expressed as a percentage of the wild-type C57BL/6 (control group) mean MFI.

### *In-situ* videomicroscopy analyses of lymphocyte interactions with PP-HEV

Naïve B cells or CD4^+^ T cells were isolated from wild-type and CD22^−/−^ or SHP-1^+/−^ splenocytes using negative selection kits according to the manufacturer’s instructions. For each experiment, the quality of the isolation was checked by flow cytometry and consisted of ≥98% cells of interest. Lymphocytes were labeled with 2.5 µM CMFDA or CMTPX for 10 minutes at 37°C in RPMI without FBS, then washed with RPMI containing 10% FBS. In some experiments, labeled B or CD4^+^ T cells were incubated with the anti-mouse α4β7 antibody DATK32 (50 µg/mL) in RPMI containing 10% FBS for 45 minutes at 37°C. Cells were washed three times to remove unbound antibodies. Prior to injection into recipient mice, cells were counted by flow cytometry using absolute counting beads. Recipient mice were anesthetized via intraperitoneal injection of ketamine and xylasine. One individual Peyer’s patch of the small intestine was exteriorized and positioned for epifluorescence microscopy and video recording. Same numbers (5-10×10^6^ cells) of wild-type, CD22^−/−^, SHP-1^+/−^ B or CD4^+^ T cells were transferred into anesthetized recipients. The interactions of fluorescent cells with PP-HEVs were recorded for 30-40 seconds at 40 frames per second of 25 msec exposure time. All fluorescent cells entering HEVs during the recording time were analyzed on a frame-to-frame basis for the entire duration of the movie. Cells passing HEVs <1 s with no significant interaction or with a velocity > 300 µm/s were considered noninteractive and called flyer; cells which started binding to HEVs briefly for <1s before getting releasing and non-interactive were considered as brief rollers. Cells binding to HEVs >1s were considered rollers. At the end of the recording, rollers attached to HEVs with a static binding of > 2s were considered arresters. To assess the mean rolling velocity of each cell, we tracked manually each cells in order to define the precise rolling distance from the frame of its first interaction (f_first_) to the first frame of its definitive static binding (f_last_). The rolling time was (f_last_ – f_first_ + 1) x 25 ms.

### Short-term homing assay

Donor splenocytes were isolated and were labeled with CTV or CFSE, or a combination of CTV and CFSE, at concentrations optimized for a bar coding system with 2–4 donor populations total. The different donor groups received a different label in every experiment to rule out an effect of the dye. Equal numbers (25–50×10^6^ cells) of donor cells were transferred into wild-type recipient mice by injection into the tail vein. After 1.5 hour, lymphocytes from the PLNs (inguinal, axillary and brachial lymph nodes) and PPs of recipient mice were isolated, stained for flow cytometry with anti-CD3, anti-CD4, anti-CD19 and anti-IgD (identified above) to identify CD 4^+^ T cell and mature B cell subsets. The Aqua dye was used for live/dead staining and counting beads were added to calculate the total numbers of cells. The homing of live CD19^+^ IgD^+^ B cells and live CD3ε^+^ CD4^+^ T cells from each donor was evaluated. For both B and CD4^+^ T cells, the efficiency of cell homing to each organ was calculated as a ratio: the number of cells found in each organ was divided by the number of cells injected (input). Then, results were presented as a percentage of the wild-type C57BL/6 (control group) mean ratio.

### Endocytosis assay

Splenocytes were stained with fluorescently labeled antibodies against CD4, CD3ε, CD19, IgD and either CD71(TfR1), CD29 (β_2_), CD49d (α_4_), β7, or IgM or the appropriate isotype control. The Fc receptors were blocked anti-CD16/32 prior to the staining. For each sample, staining at 4°C with Phycoerythrin (PE)-conjugated RI7217 (anti-Transferrin Receptor 1, TfR1), HMβ1-1 (anti-β_1_), FIB504 (anti-β_7_) and R6-60.2 (anti-IgM), or the matching PE-conjugated isotype control antibodies allowed calculation for extra-cellular levels of TfR1, β_1,_ α_L_, β_7_, and IgM levels. In parallel, staining of each sample with pHrodo-Red-conjugated Transferrin (Tf), HMβ1-1, R6-60.2 (anti-IgM), and FIB504 (anti-β_7_) antibodies at either 4°C (no endocytosis) or 37°C (endocytosis) was used to calculate endocytosis levels. The same stainings were also performed in the presence of 100µM Primaquine (PQ). For each antigen and each experiment, the Relative Endocytosis Ratio (RER) was calculated by normalizing endocytosis levels to extra-cellular levels, i.e.:

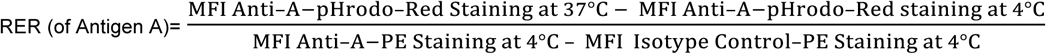

### Real-time quantitative RT-PCR

Naïve B cells were isolated from wild-type and CD22^−/−^ splenocytes using a mouse B cell selection kit according to the manufacturer’s instructions. Total RNA was isolated from purified B cells using the RNeasy mini kit. Analysis of total RNA concentration and integrity was assessed using Nanodrop ND-1000 spectrophotometry. Equal amounts of each total RNA samples were converted into cDNA using the High Capacity cDNA Reverse Transcription Kit according to the manufacturer’s instructions. Quantitative RT-PCR analysis of genes of interest was performed using the PowerSYBR**®** Green PCR Master Mix on a ABI PRISM® 7900HT (Applied Biosystems). PCR reaction mixtures consisted of: 1uL of a 1:5 dilution of template cDNA, 200nM of each primer, and 1X PowerSYBR**®** Green PCR Master Mix in a final volume of 10uL. RT-qPCR amplication was conducted using an initial step of 5 min at 45°C, then 5 min at 95°C, followed by 40 cycles of denaturation at 95°C for 30 sec, primer annealing at 60°C for 30 sec, and extension at 72°C for 30 sec. Melting curve analysis was used to assess the purity of the amplified bands. Sequences for the primer pairs used are as follows: β_1_(DeNucci et al., 2010), 5’-AATGCCAAATCTTGCGGAGAA-3’ (forward) and 5’-TCTAAATCATCACATCGTGCAGAAGTA-3’ (reverse); β_2_(DeNucci et al., 2010) 5’-GATAACATGTACAAGAGGAGCAATGAGT-3’ (forward) and 5’-CGCAAAGATGGGCTGGAT-3’ (reverse); α_L_(DeNucci et al., 2010), 5’-AGGTTGACCTGATCCACGAG-3’ (forward) and 5’-CAGGTTCCGTTTGAAGAAGC-3’ (reverse); β_7_(Eun et al., 2007), 5′-CTATCCTCCCTTTCTCTATCAG-3′ (forward) and 5′-GTCTAGGTAGCGCTTGCTTAC-3′ (reverse); β-actin (pre-designed primers from Qiagen); HPRT(Eun et al., 2007), 5’-AGGGATTTGAATCACGTTTG-3’ (forward) and 5’-TTTACTGGCAACATCAACAG-3’ (reverse). We used the dCT (CT) method to calculate the relative gene mRNA expression of α_L_, β_1_, β_2_, and β_7_ using β-actin and HPRT as housekeeping genes.

### CTB immunization studies

Adults C57BL/6 or CD22^−/−^ mice were immunized with 10 µg Cholera Toxin B diluted in 100 µL PBS for oral gavage (Oral route), 10 µL PBS for intra nasal via pipetting in the nostrils (Nasal route), or 50 µL PBS for injection into the mouse hamstring (Muscular route). Mice were sacrificed two weeks post immunizations. Serum was collected for ELISA studies, and segments of small intestine used in *ex-vivo* cultures.

### Rotavirus infection studies

Five days old C57BL/6 or CD22^−/−^ pups were orally gavaged with 10^4^ diarrhea dose 50% (DD_50)_ of wild-type murine rotavirus, strain EW, diluted in M199 medium. Pups were then checked daily for diarrhea by gentle abdominal pressure. Fecal samples from each individual mouse were collected in PBS with Calcium/Magnesium for enzyme-linked immunosorbent assays (ELISAs). In some studies, mice were sacrificed either at day 12 p.i. to collect the serum for ELISA studies, mesenteric lymph nodes (MLN), small intestine lamina propria (SI LP) and spleen for ELISPOT analysis, and segments of SI for *ex-vivo* cultures.

### Detection of viral antigen and virus specific IgA and IgG by ELISA

For detection of viral antigen in fecal samples(Franco and Greenberg, 1995b), 96-well ELISA plates were coated with guinea pig anti-rotavirus hyperimmune serum (1:4,000 in PBS) and incubated at 37°C for 4 h. The plates were then blocked with 5% non-fat milk in PBS (Blotto) at 37°C for 2 h. Suspended fecal samples were diluted 1:20 in 1% Blotto, added to the plates, and incubated at 37°C for 1 h. The plates were washed three times with PBS containing 0.05% Tween 20 (PBS-T). Rabbit anti-rotavirus hyperimmune serum (1:3,000 in 1% Blotto) was added to the plates for 1 h at 37°C, and washed three times. Horseradish peroxidase-conjugated goat anti-rabbit IgG (1:30,000 in 1% Blotto) was added to the plates and incubated for 1 h at 37°C. TMB substrate was added after four washes in PBS-T, the plates were developed for 10 min at room temperature, the reaction was stopped by the addition of 0.16 M sulfuric acid, and the OD450 read with a plate reader. Data are presented as OD450-like numbers after normalization to stools weight.

For the detection of virus-specific fecal and serum IgA and IgG, plates were coated with rabbit anti-rotavirus hyperimmune serum and blocked as for the antigen detection ELISA, and then incubated with a dilution of rotavirus stock (1:5 in 1% Blotto) overnight at 4°C. After three washes with PBS-T, stool samples (1:20 in Blotto) or serial dilutions of serum in 1% Blotto were added to the plates. After 2 h of incubation at 37°C, the plates were washed three times in PBS-T and peroxidase-conjugated anti-mouse IgA or IgG (diluted 1:1,000 and 1:20,000 respectively in 1% Blotto) were added for 1 h at 37°C. Plates were washed and developed as described above. For the detection of Total fecal and serum IgA or IgG, plates were coated with anti-mouse IgA, IgG, IgM polyconal antibodies (2 µg/mL in PBS) before using the same protocol as above.

### Detection of CTB-specific IgA and IgG by ELISA

ELISA plates were coated with CTB (2 µg/mL) overnight, blocked with in PBS containing 5% BSA, then incubated with serial dilutions of serum or *ex-vivo* SI culture samples diluted in PBS containing 2% BSA for 2 h. After four washes with PBS-T, plates were peroxidase-conjugated anti-mouse IgA or IgG (diluted 1:1,000 and 1:10,000 respectively in PBS containing 2% BSA) were added for 1 h at 37°C. TMB substrate was added after four washes in PBS-T, the plates were developed for 5-10 min at room temperature, the reaction was stopped by the addition of 0.16 M sulfuric acid, and the OD450 read with a plate reader. For the detection of serum and *ex-vivo* SI culture samples IgA or IgG, plates were coated with anti-mouse IgA, IgG, IgM polyconal antibodies (2 µg/mL in PBS) before using the same protocol as above.

### Small intestine fragment cultures

Open segments (1 cm long) of duodenums were weighted and placed in 24-well plates on elevated mesh filter and cultured for three days at 37°C in complete RPMI medium in hyper oxygenated conditions(Feng et al., 2006). At the end of the incubation period, intestinal IgA and IgG were measured by ELISA in the supernatants as described above.

### Quantitation of virus-specific antibody secreting cells (ASCs) by Elispot assay

Multiscreen 96-well plates (MAIPS4510, Millipore) were coated with purified rotavirus double-layer particle overnight at 4°C, washed, and blocked with RPMI medium containing 10% FCS for 1 h at 37°C. Serial 10-fold dilutions of spleen, PP, or SI LP cells were added to the plates, and incubated overnight at 37°C. After six washes, peroxidase-conjugated anti-mouse IgA or IgG were added for 1 h at 37°C. Plates were washed six times and developed with AEC substrate. To quantity non specific ASCs, plates were coated with anti-mouse IgA, IgG, IgM polyclonal antibodies (1:250) before using the same protocol as above.

---

**Figure S1.**
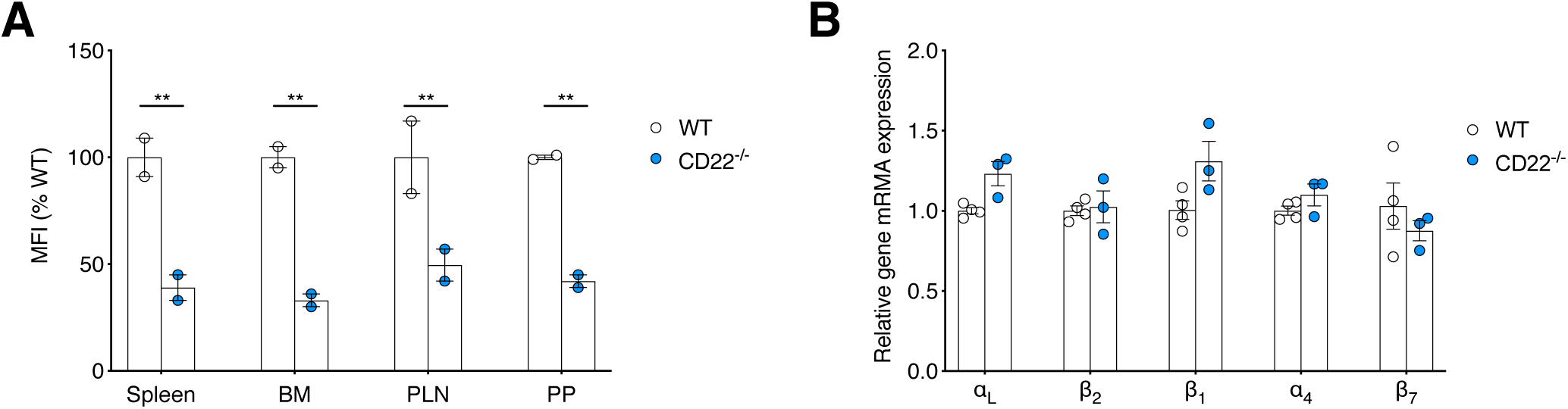
Systemic yet non-transcriptional CD22-dependent regulation of α_4_β_7_ on B cells **(A)** Flow cytometry of wild-type (WT) or CD22-deficient (CD22^−/−^) live CD19^+^ IgD^+^ B cells isolated from the spleen, the bone marrow (BM), the peripheral lymph node (PLN), and the Peyer’s patches (PP) and stained with antibodies against α_4_β_7_ or isotype-matched control. Shown are pooled data (mean ± SEM) from n=2 independent experiments with two animal per group total. The median fluorescence intensity (MFI) of the integrin staining was expressed as a percentage of the WT B cells group mean. ** P<0.01 (Two-way ANOVA with Sidak’s multiple comparison test). **(B)** mRNA expression of α_L_,β_2_, β_1_, α4, and β_7_ integrins in B cells purified from spleens of WT and CD22^−/−^ mice, normalized to that of actin and HPRT used as housekeeping genes. Shown are pooled data from two independent experiments with three animals per group.

**Figure S2.**
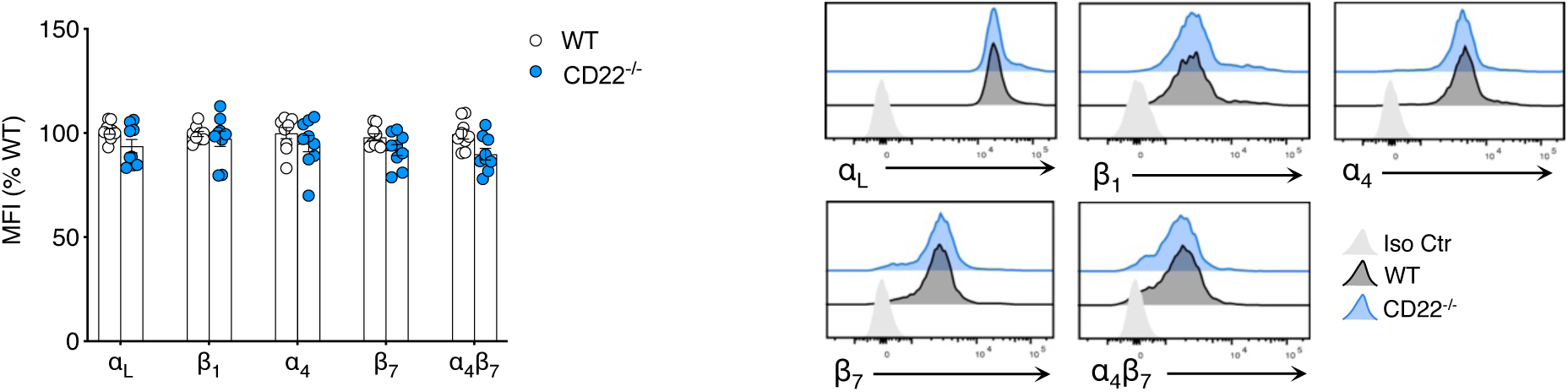
CD22^−/−^ T cells display normal cell surface levels of α_4_β_7_. Flow cytometry of WT or CD22^−/−^ live CD3^+^ CD4^+^ T cells isolated from spleens and stained with antibodies against the integrins α_L_, β_1_, α_4_, β_7_, or α_4_β_7_. Shown are pooled data (mean ± SEM) from n=3 independent experiments with 7 animals per group total presented as in **Figure 1**. Representative histogram overlays are shown.

**Figure S3.**
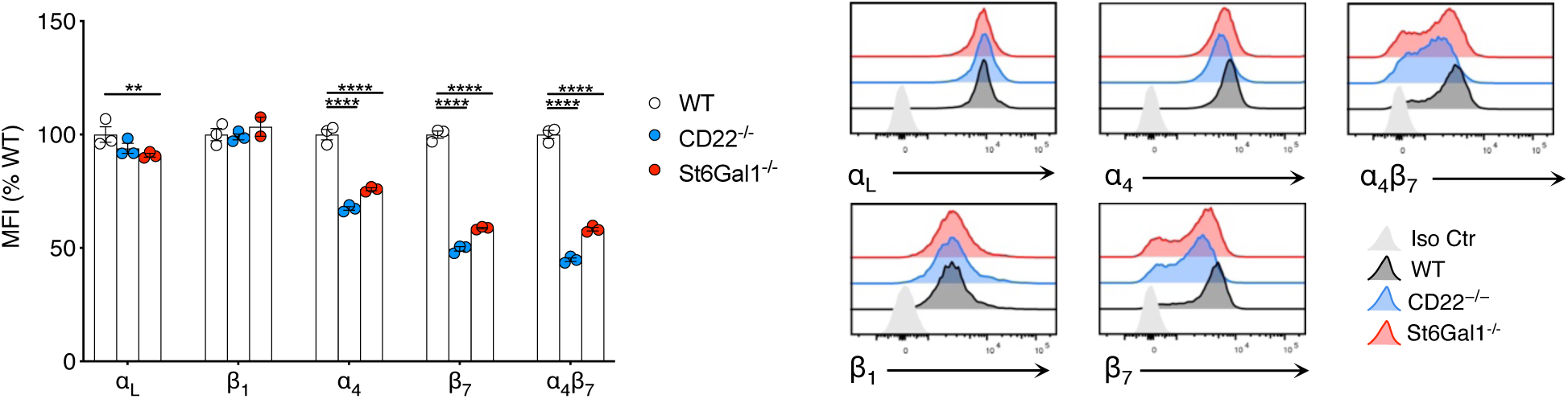
B cell expression of CD22-binding carbohydrate control α_4_β_7_ expression. Flow cytometry of WT, CD22^−/−^, or St6Gal1^−/−^ naïve B cell isolated from spleen and stained with antibodies against the integrins α_L_, β_1_, α_4_, β_7_, or α_4_β_7_. Data represent the mean ± SEM of one representative experiment with n=3 mice per group presented as in **Figure 1**. Representative histogram overlays are shown. Groups were compared by one way ANOVA with Dunnett’s multiple comparisons test. **P<0.01; and \*\*\*\**P* < 0.0001.

**Figure S4.**
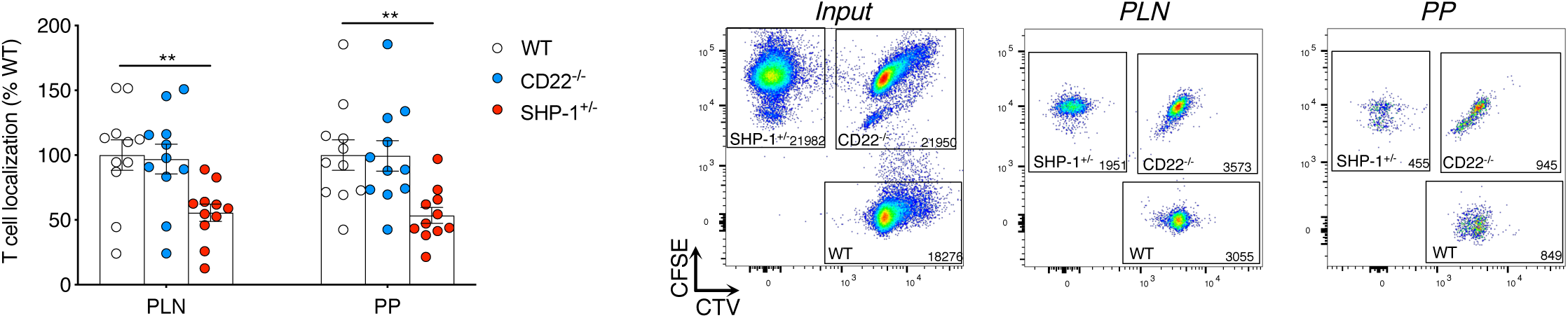
Localization of WT, CD22^−/−^, and SHP-1^+/−^ T cells in PLN and PP after short-term homing assays. WT, CD22^−/−^, or SHP-1^+/–^ splenocytes labeled with CFSE, or CellTracker Violet (CTV) or both were injected i.v. into a recipient WT mouse. PLN, MLN or PP cells isolated from the recipient were stained with anti-CD3 and anti-CD4 for quantification of short-term (90 min) homing of CD4 T cells. For each donor and each organ, the number of isolated CD4 T cells (Output) was normalized to the number of injected CD4 T cells (Input) to yield a T cell localization ratio. Shown is the mean ± SEM from three independent experiments with n=11 mice per group total. Representative dot plots gated in live CD3^+^ CD4^+^ T cells are shown, including the number of cells within each gate. **P<0.01 (Two-way ANOVA with Sidak’s multiple comparison test).

**Figure S5.**
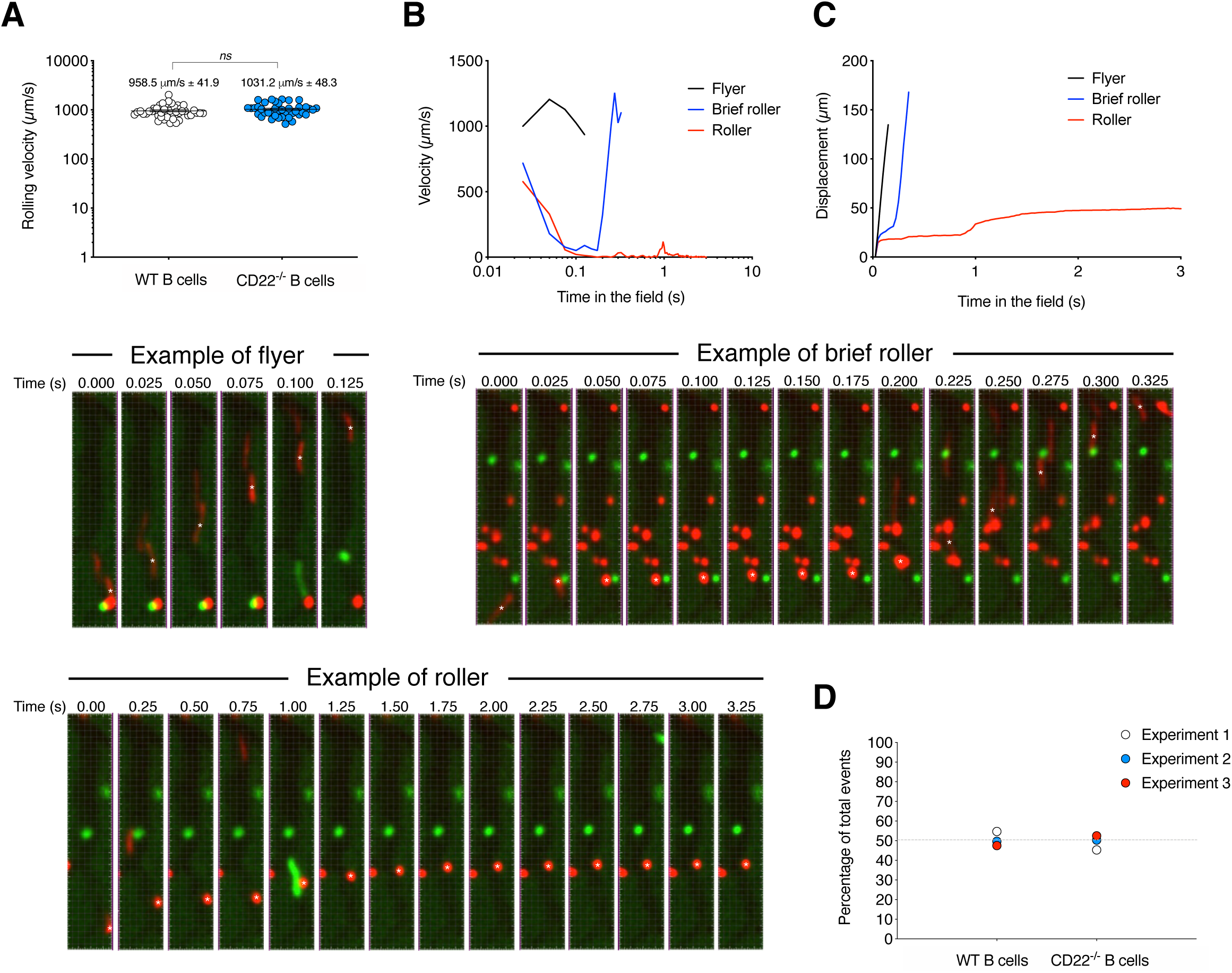
Definition of flyer, brief roller and roller cells visualized by *in situ* videomicroscopy of Peyer’s patches. **(A)** The mean velocity of wild-type (WT) and CD22-deficient (CD22^−/−^) B cells free flowing through the vessels without any interactions (namely flyer) was calculated and shown for each individual cell. Shown are pooled results (∼ 50 cells per group) analyzed from 3–4 representative HEVs and 3 independent experiments. **(B-C)** The instant velocity **(B)** and displacement **(C)** of a representative free flowing cell (Flyer), of one that interacts very briefly (<1s) with the HEVs (namely Brief roller), or one that interacts and rolls on the HEVs for >1sec (namely Roller) is shown together with frame-per-frame tracking of the cells (identified with *). **(D)** In three independent *in situ* experiments with 1:1 ratio of WT B cells donor versus CD22^−/−^ B cells donor, the total number of events (i.e. flyer, brief roller, or roller) was counted for each donor in 3–4 representative HEVs for the total duration of the movie (∼ 250–300 total cells analyzed per group). The percentage of WT and CD22^−/−^ B cells experiment per experiment is shown.

**Figure S6.**
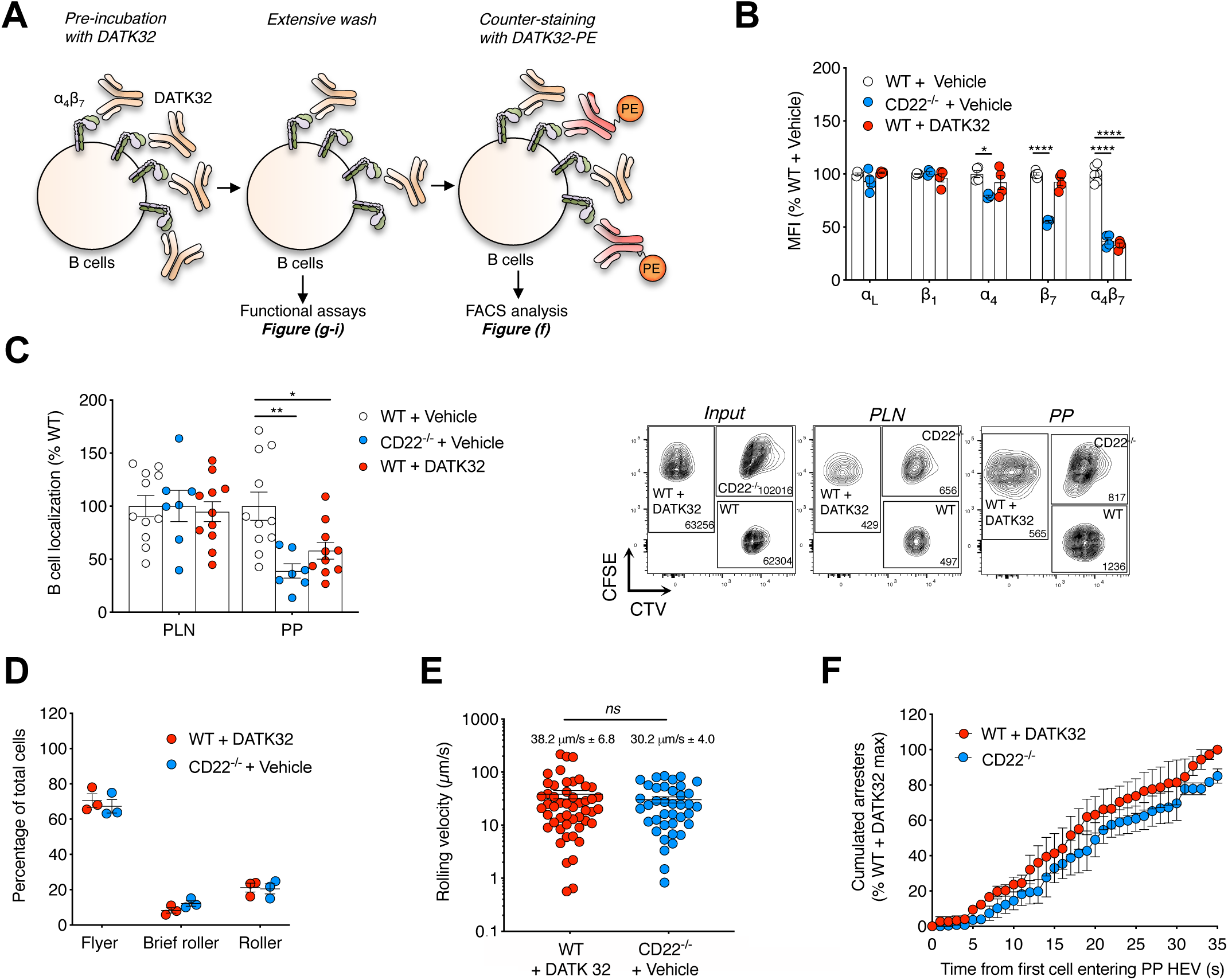
Experimental reduction of α_4_β_7_ on WT B cells mimics the defective CD22-deficient B cell homing to PP. **(A)** WT B cells were pre-incubated with inhibitory anti-α_4_β_7_ Ab DATK32 and washed extensively. DATK32-pretreated B cells were either counter-stained with Phyco-erythrin(PE)-conjugated DATK32 for flow cytometry analyses (B), or use in functional assays (C-F) **(B)** Flow cytometry of WT + DATK32 vs. WT + Vehicle B cells stained for α_L_, β_1_, α_4_, β_7_, or α_4_β_7_ presented as in **Figure 1**. Shown are pooled data (mean ± SEM) from n=2 independent experiments with 4 animals per group total. **(C)** Localization of WT, CD22^−/−^, and WT + DATK32 B cells in PLN and PP after homing assays analyzed and presented as in **Figure 4**. Shown are pooled data (mean ± SEM) of n=3 experiments with 11 mice per group total. Representative dot plots gated in live naïve B cells are also shown including the number of cells within each gate. **(D-F)** *In situ* videomicroscopy analyses of WT B cells + DATK32 vs. CD22^−/−^ B cells interactions with PP-HEVs analyzed and presented as in **Figure 4**. Data represent the mean ± SEM of three independent experiments (D,F) and representative cells from all 3 experiments (E). Groups were compared by one way ANOVA with Dunnett’s multiple comparisons test (B,C), unpaired two-tailed Student’s *t*-test (D,E), and paired two-tailed Student’s t-test (F). \**P* < 0.05, \*\**P* < 0.01, \*\*\*\**P* < 0.0001. NS, not significant.

**Figure S7.**
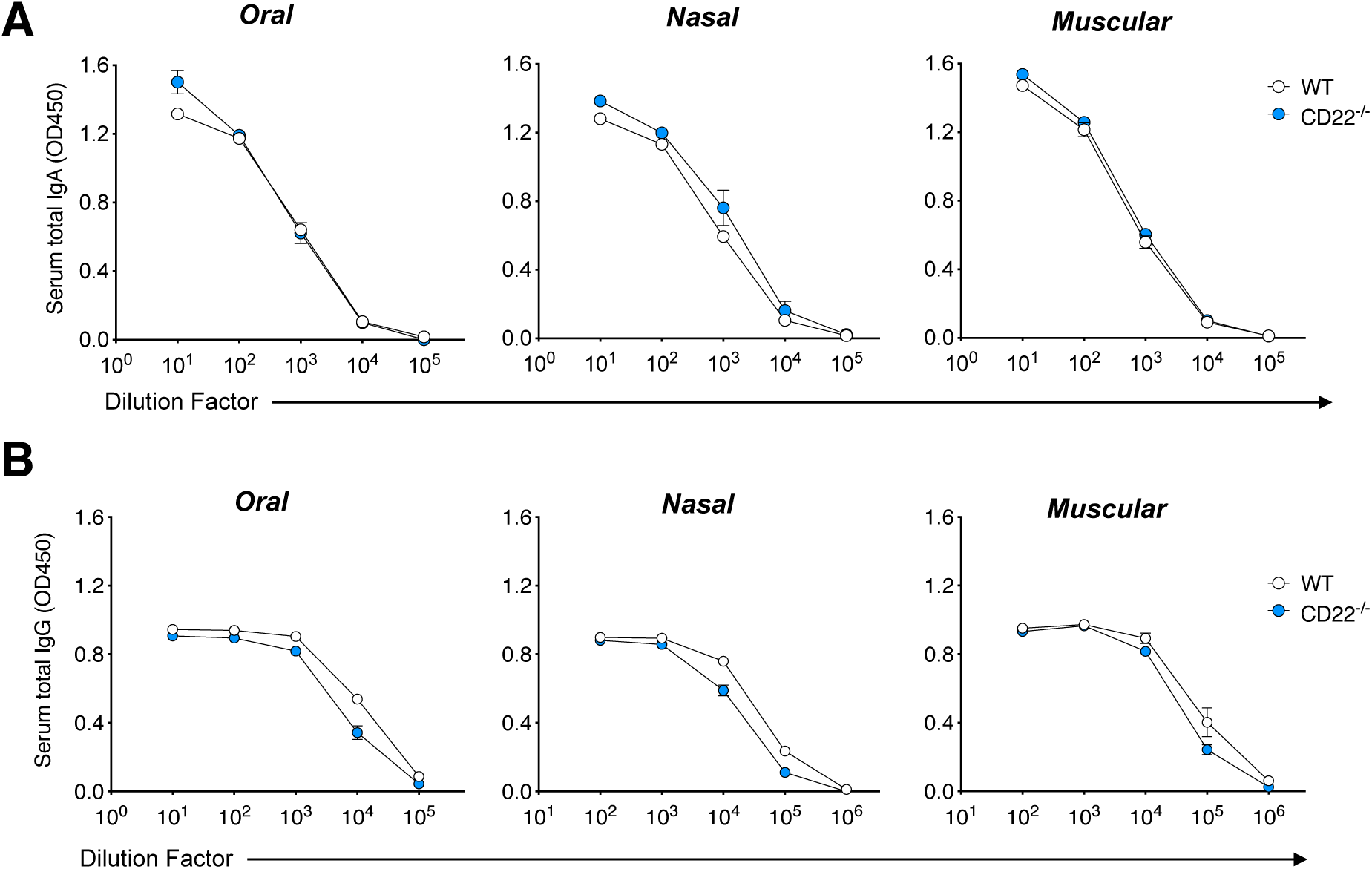
Similar total IgA and IgG serum titers in wild-type and CD22-deficient mice after CTB immunizations. Cohorts of WT or CD22^−/−^ mice were immunized with Cholera Toxin B (CTB) via the Oral, intra-nasal (Nasal), or intra-muscular (Muscular) routes for two weeks. The serum levels of total IgA **(A)** or IgG **(B)** were measured by ELISA and expressed as net OD450. Shown are the mean ± SEM with n=3 mice per group from one out of two independent experiments with similar results.

**Figure S8.**
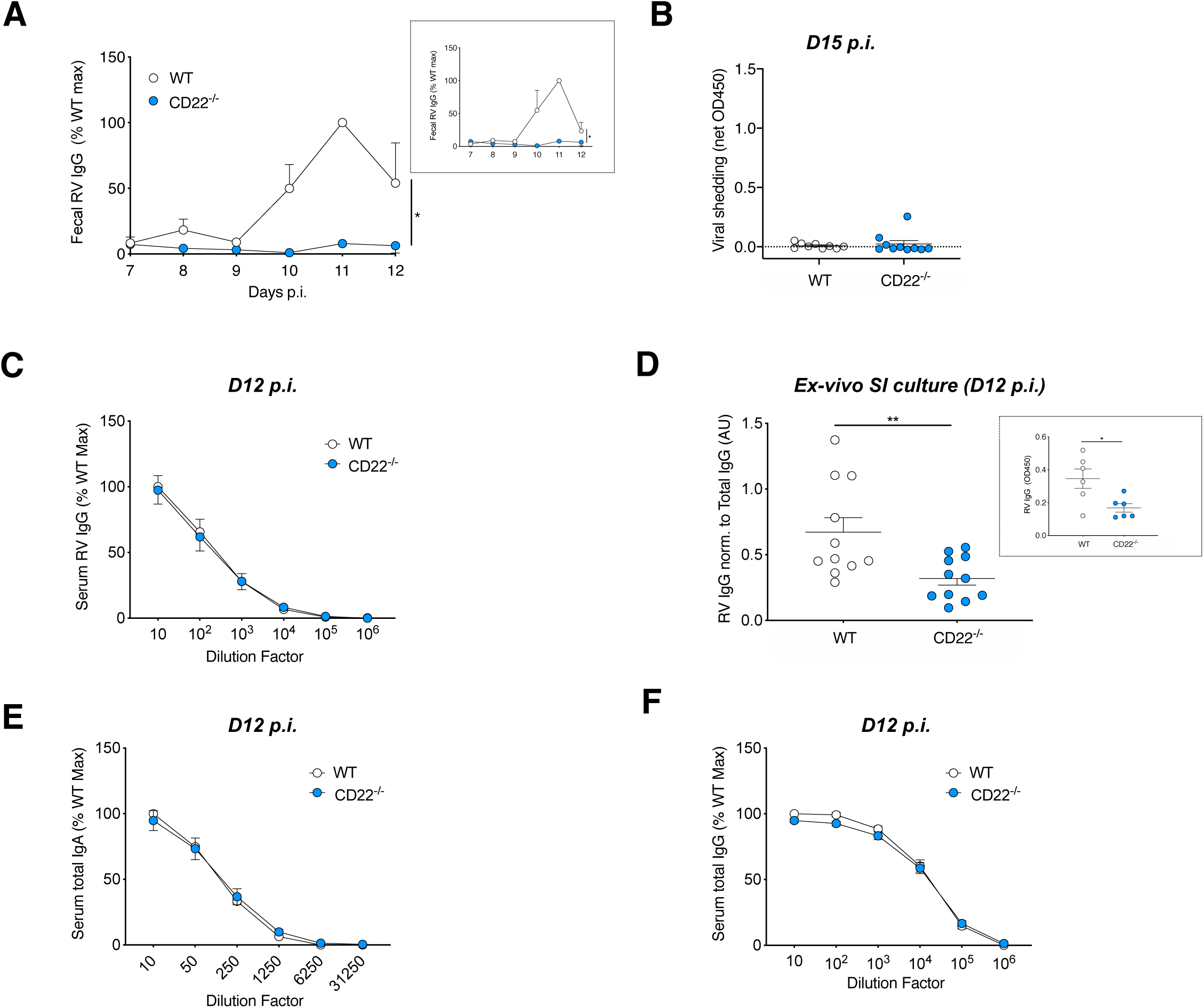
Reduced intestinal RV IgG response to rotavirus (RV) infection in CD22-deficient animals, but normal systemic total IgA and IgG titers. **(A-B)** Five days old WT or CD22^−/−^ pups were orally gavaged with the RV strain EW. **(A)** The production of RV-specific IgG and total IgG in fecal samples was measured by ELISA up to twelve days post-infection. For each individual animal within one experiment, the ratio of RV IgG to total IgG levels was expressed as a percentage of the maximal ratio in the WT group. Shown are pooled data (mean ± SEM) from n=2 independent experiments with 5-10 animals per group total. Boxed inset: ratios of RV IgG to total IgG shown as arbitrary unit (AU) in one representative experiment. **(B)** Fecal RV antigen shedding at day 15 p.i. as measured by ELISA and expressed as net OD450, normalized to the sample weight. **(C-F)** In separate experiments, five days old WT and CD22^−/−^ pups were orally inoculated with RV EW and sacrificed at day twelve post-infection. Shown in panels (C-F) are pooled data (mean ± SEM) from n=2 independent experiments with 10-11 mice per group **(C)** The serum levels of RV-specific IgG were measured by ELISA. For each experiment, data are expressed as a percentage of the maximal net OD450 measured in the WT group at the lowest serum dilution. **(D)** Ratios of RV IgG to total IgG in *ex-vivo* SI segments cultures (AU). Boxed inset: RV IgG titers shown as net OD450 in one out of two experiments. **(E-F)** The serum levels of total IgA **(E)** or IgG **(F)** were measured by ELISA and presented as in panel (C). Groups were compared by paired one-tailed Student’s t-test (A), and unpaired two-tailed Student’s t-test (D).*P<0.05; **P<0.01.

**Figure S9.**
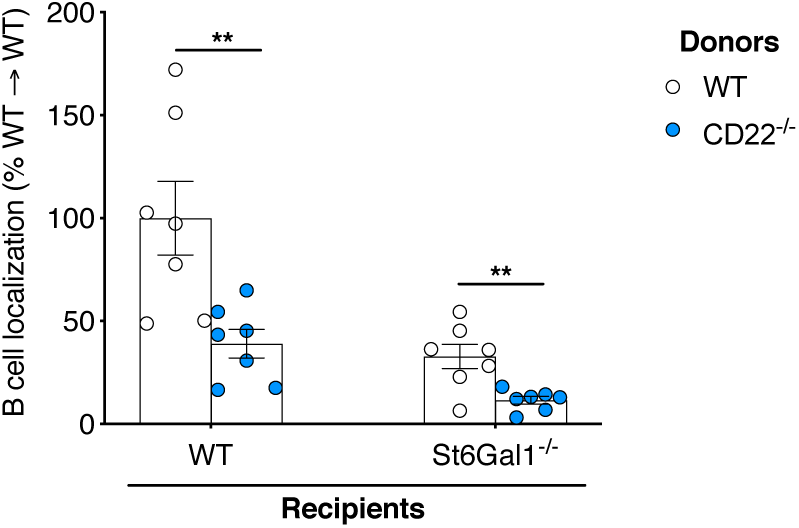
Defective homing of CD22^−/−^ B cells in ligand-deficient recipient mice. Localization of WT and CD22^−/−^ B cells in the PPs of wild-type (WT) or ligand-deficient (St6Gal1^−/−^) mice after short-term (1.5 hr) homing assays designed as in **Figure 4A**. For each donor, the number of B cells isolated (Output) was normalized to the number of injected B cells (Input), and shown as a percentage of the WT→WT group mean. Shown are pooled data (mean ± SEM) from three experiments with n=7 mice per group total. ** P<0.01 (Student‘s t-test).

**Figure S10.**
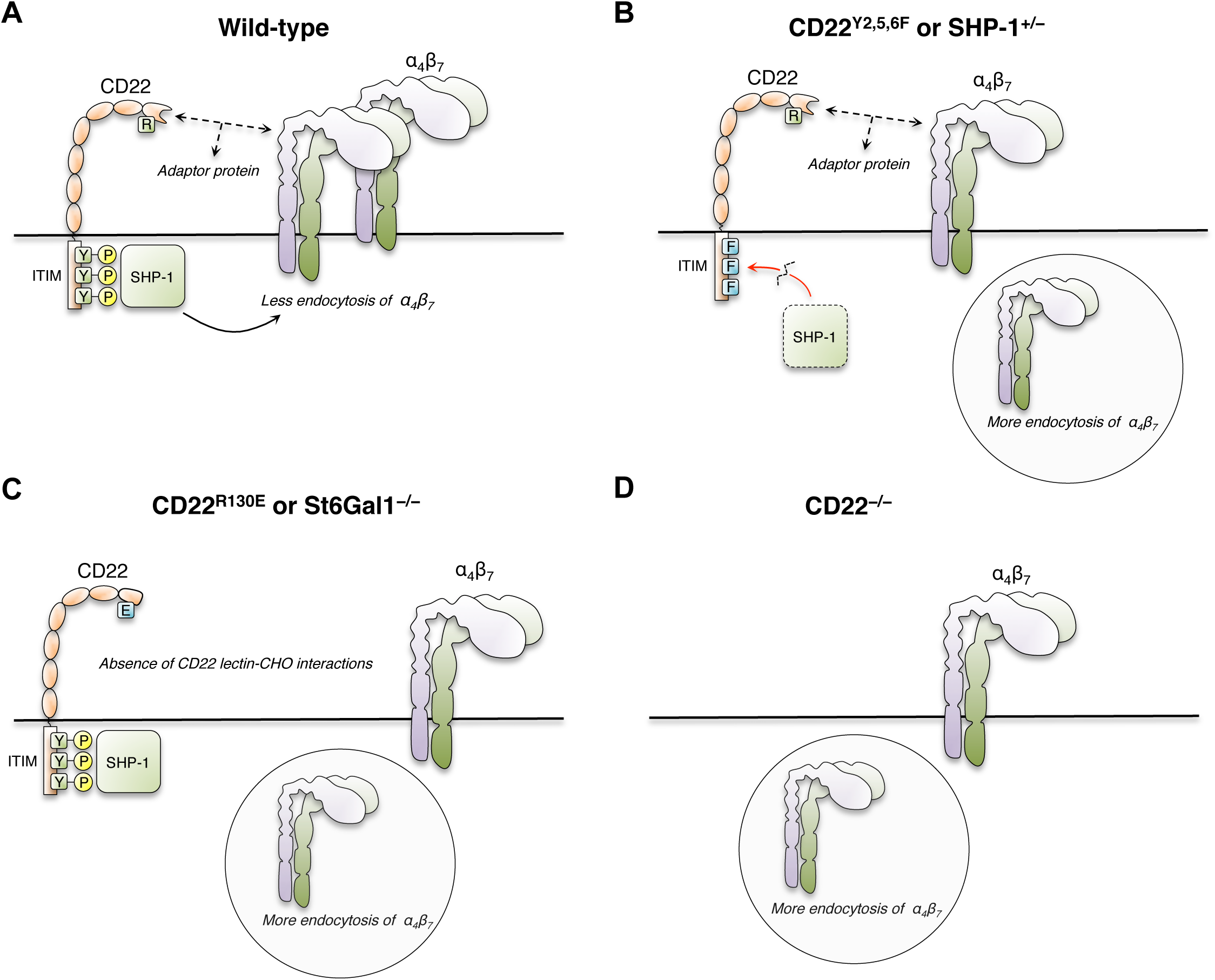
Our working hypothesis. **(A)** In wild-type B cells, SHP-1 is targeted to the cell surface by the CD22 ITIM and lectin domains to inhibit endocytosis of the integrin β_7_ **(B)** In transgenic animals with multiple mutations in SHP-1-binding ITIM sequences of CD22 (CD22^Y2,5,6,F^ knock-in) or in SHP-1 haploinsufficient animals (SHP-1^+/−^), the CD22 lectin activity remains intact, but the defective targeting of SHP-1 to the CD22 ITIM domains results in increased β_7_ endocytosis. **(C)** In transgenic animals with a single point mutation in the CD22 lectin binding domain (CD22^R130E^ knock-in) or in animals deficient in the enzyme (St6Gal1) that forms the CD22 ligand (St6Gal1^−/−^), the recruitment of SHP-1 to the CD22 ITIMs remains functional, but absence of lectin-carbohydrate (CHO) interactions can not bring ITIM-bound SHP-1 phosphatase in close proximity to target substrates, resulting in higher β_7_ endocytosis. **(D)** The absence of CD22 (CD22^−/−^) is a combination of the scenarios (B) and (C), also resulting in accelerated β_7_ endocytosis.

**Supplemental Video 1.**
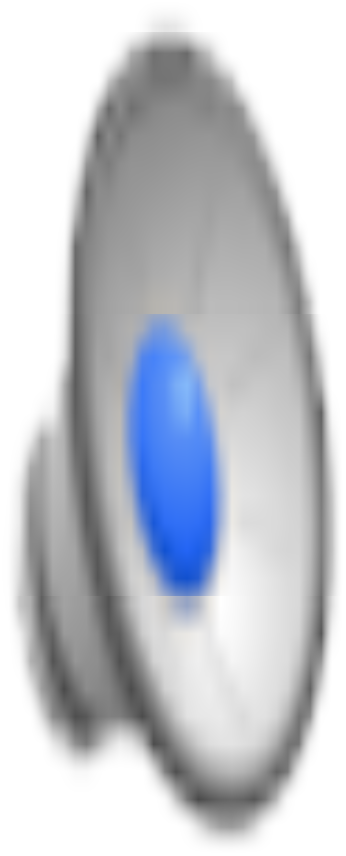
Example of flyer visualized by *in situ* videomicroscopy of Peyer’s patches. The same example of free flowing cell shown frame per frame in **Figure S5** is shown here as a movie (5 frames per second, 0.125 sec movie). Scale bar, 10*µ*m.

**Supplemental Video 2.**
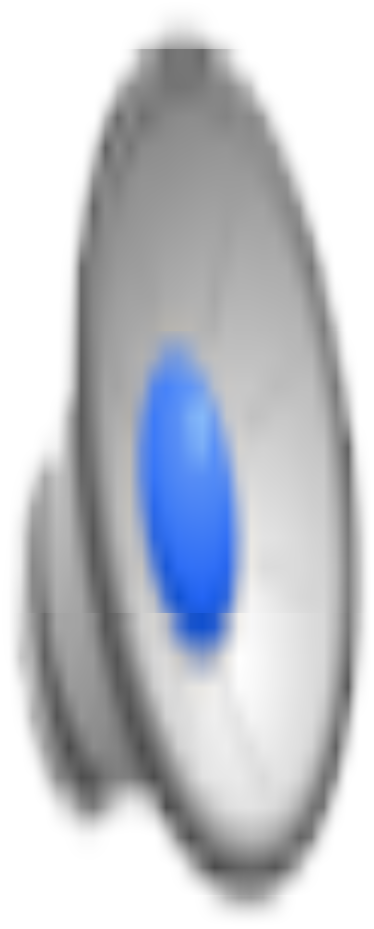
Example of brief roller visualized by *in situ* videomicroscopy of Peyer’s patches. The same example of brief roller shown frame per frame in **Figure S5** is shown here as a movie (5 frames per second, 0.325 sec movie). Scale bar, 10*µ*m.

**Supplemental Video 3.**
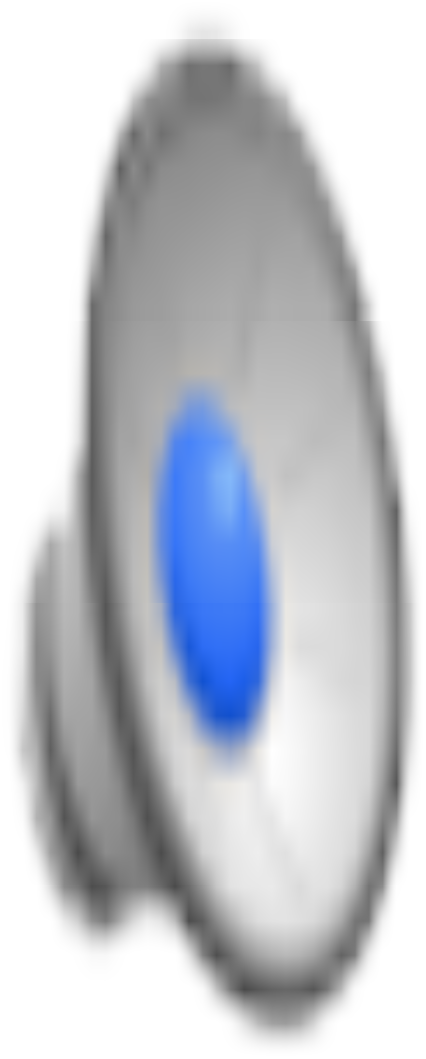
Example of roller visualized by *in situ* videomicroscopy of Peyer’s patches. The same example of roller shown every 10 frames in **Figure S5** is shown here as a movie (20 frames per second, 3.275 sec movie). Scale bar, 10*µ*m.

**Supplemental Video 4.**
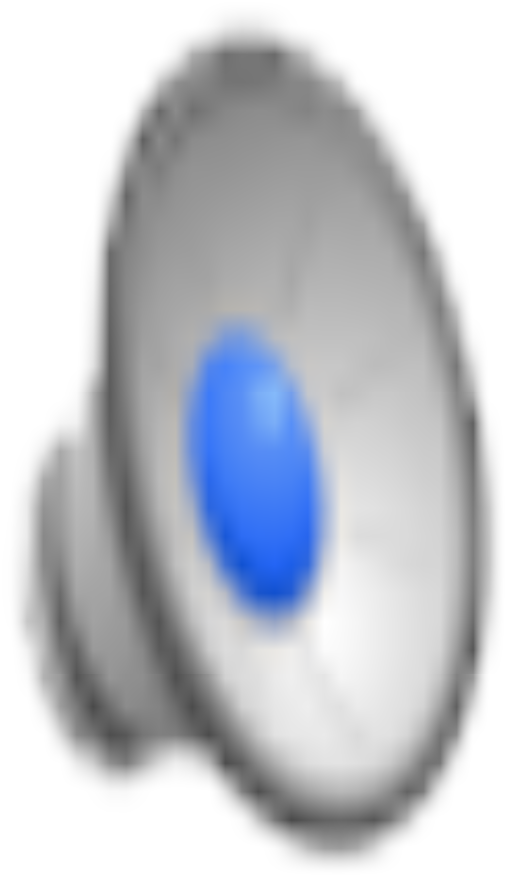
*In situ* videomicroscopy reveals defective arrest of SHP-1^+/−^ B on PP HEVs (Related to **Figure 4**). Representative real-time videomicroscopy experiment showing purified wild-type (in green) and SHP-1^+/−^ B cells (in red) interacting with PP HEVs. (40 frames per second, 32.23 sec movie). Scale bar, 50*µ*m.

**Supplemental Video 5.**
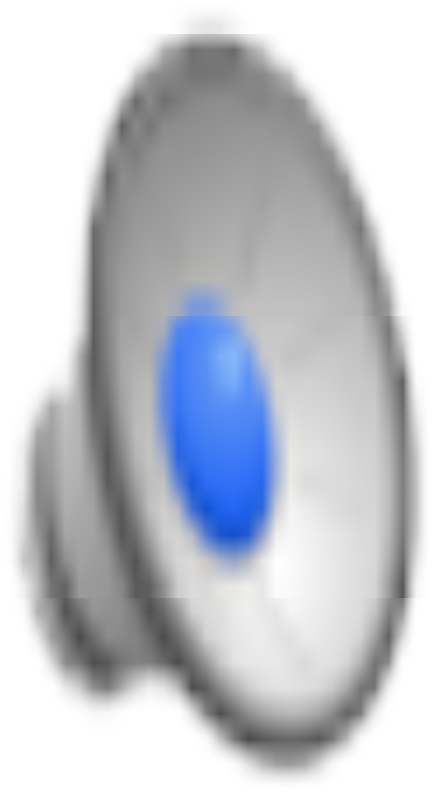
*In situ* videomicroscopy reveals defective arrest of CD22^−/−^ B on PP HEVs (Related to **Figure 4**). Representative real-time videomicroscopy experiment showing purified wild-type (in red) and CD22^−/−^ B cells (in green) interacting with PP HEVs. (40 frames per second, 41.628 sec movie). Scale bar, 50*µ*m.

**Supplemental Video 6.**
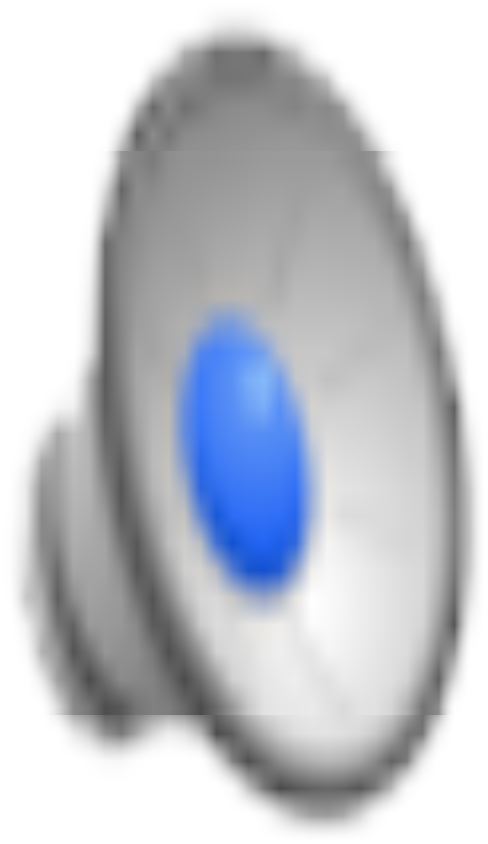
*In situ* videomicroscopy reveals increased rolling velocity of SHP-1^+/−^ B on PP HEVs (Related to **Figure 4**). Representative wild-type B cell roller (green) and SHP-1^+/−^ B cell roller (red) interacting with PP HEVs. (40 frames per second; 10.48 sec movie). Scale bar, 10*µ*m.

**Supplemental Video 7.**
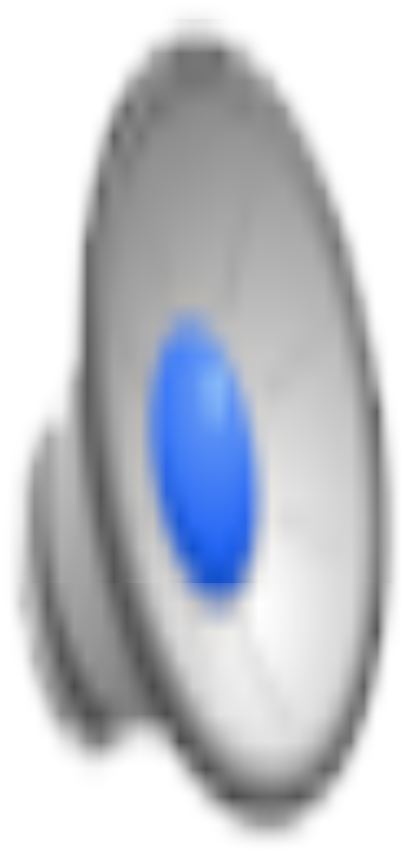
*In situ* videomicroscopy reveals increased rolling velocity of CD22^−/−^ B on PP HEVs (Related to **Figure 4**). Representative wild-type B cell roller (red) and CD22^−/−^ B cell roller (green) interacting with PP HEVs. (40 frames per second; 12,715 sec movie). Scale bar, 10*µ*m.

